# Two closely related RNA helicases, UAP56 and URH49, couple mRNA splicing and export pathways with their distinct complexes

**DOI:** 10.64898/2026.10.27.762353

**Authors:** Ken-ichi Fujita, Kotaro Harada, Yuya Ikeda, Hanae Yoshioka, Moe Yokoshi, Fumito Hori, Naoko Fujiwara, Shigeto Seno, Hideo Matsuda, Yukio Kawahara, Akila Mayeda, Seiji Masuda

## Abstract

Transcription and mRNA processing are integrated into appropriate mRNA export pathway(s) enabling fine-tuned gene expression. The closely related RNA helicases, UAP56 and URH49, form distinct complexes known as the apo-TREX and the apo-AREX complexes, respectively, which remodel into the ATP-TREX complexes, thus forming multiple mRNA export pathways. However, the mechanisms by which helicases and their complexes shape these pathways remain unclear. Here, we showed that UAP56 and URH49 play general roles in mRNA splicing ahead of export. Further analyses revealed three patterns of splicing regulation and their association with mRNA export: Type I, in which both helicases redundantly or synergistically regulate splicing within the same transcripts, associated with cooperative mRNA export; Type II, in which either helicase preferentially regulates splicing in distinct transcripts; and Type III, in which the helicases exert opposing splicing effects within the same transcripts. Type II and III regulation was associated with selective mRNA export. Moreover, we demonstrated that the distinct complex formation influenced the splicing patterns regulated by the two helicases. Our findings reveal an additional layer of transcriptome regulation mediated by UAP56, URH49, and their complexes, extending beyond their canonical roles in mRNA export.

## Introduction

The lifetime of an mRNA involves several stages leading to the expression of protein-coding genes. The pre-mRNA is transcribed and undergoes several RNA processing events: 5’-capping, splicing, and 3’-end processing, in the nucleus. Among these processes, splicing is executed by the removal of intron(s) from pre-mRNA by the spliceosome, composed of a core of U snRNA ribonucleoprotein (U snRNP) complexes, and accessory proteins(1). The spliceosome recognizes short consensus sequences in the intron(s) of pre-mRNA, at the 5’ splice site, the branchpoint sequence (BPS), and the poly-pyrimidine tract (PPT) followed by the 3’ splice site. The usage of splice sites is variable and regulated by factors associated with the spliceosome to generate a diverse array of alternatively spliced isoforms that contribute to a wide range of biological processes(2,3). Following mRNA processing, the mature mRNA is exported to the cytoplasm for protein translation. mRNA export plays a crucial role in gene expression and is coordinately processed with mRNA processing through the appropriate assembly and remodeling of the mRNA-protein complexes (1).

Integration from transcription to mRNA export is facilitated by the evolutionarily conserved ATP-dependent multi-subunit <u>TR</u>anscription-<u>EX</u>port (TREX) complex. The human TREX complex consists of the THO subcomplex, several affiliated proteins, and one of two DEAD-box RNA helicases, UAP56 (also known as DDX39B) or its closely related paralogue, URH49 (also known as DDX39A, Fig.1a)(4,5). The TREX complex components are recruited onto transcribing RNA polymerase II (Pol II)(4) and loaded onto spliced-mRNA in a splicing-dependent manner(6). We previously found that the TREX complex exists in two states: the apo-TREX complex containing UAP56 and the THO subcomplex, and the ATP-TREX (UAP56) complex, which contains all known TREX components through ATP binding to UAP56, facilitating complex remodeling (Fig.1a)(5). The formation of the apo-TREX complex is crucial for the recognition of target mRNAs for export(7). Furthermore, subsequent remodeling from the apo-TREX to the ATP-TREX (UAP56) complex triggers mRNA export, because the ATP-TREX (UAP56) complex components, ALYREF, CHTOP, and THOC5 serve as a platform for the mRNA exporter NXF1-NXT1 heterodimer(8–10). More recently, structural and biochemical studies have revealed interactions between UAP56 and the TREX-2 complex, suggesting a role in mRNP remodeling and its transfer to NXF1–NXT1(11–13).

**Fig.1.**
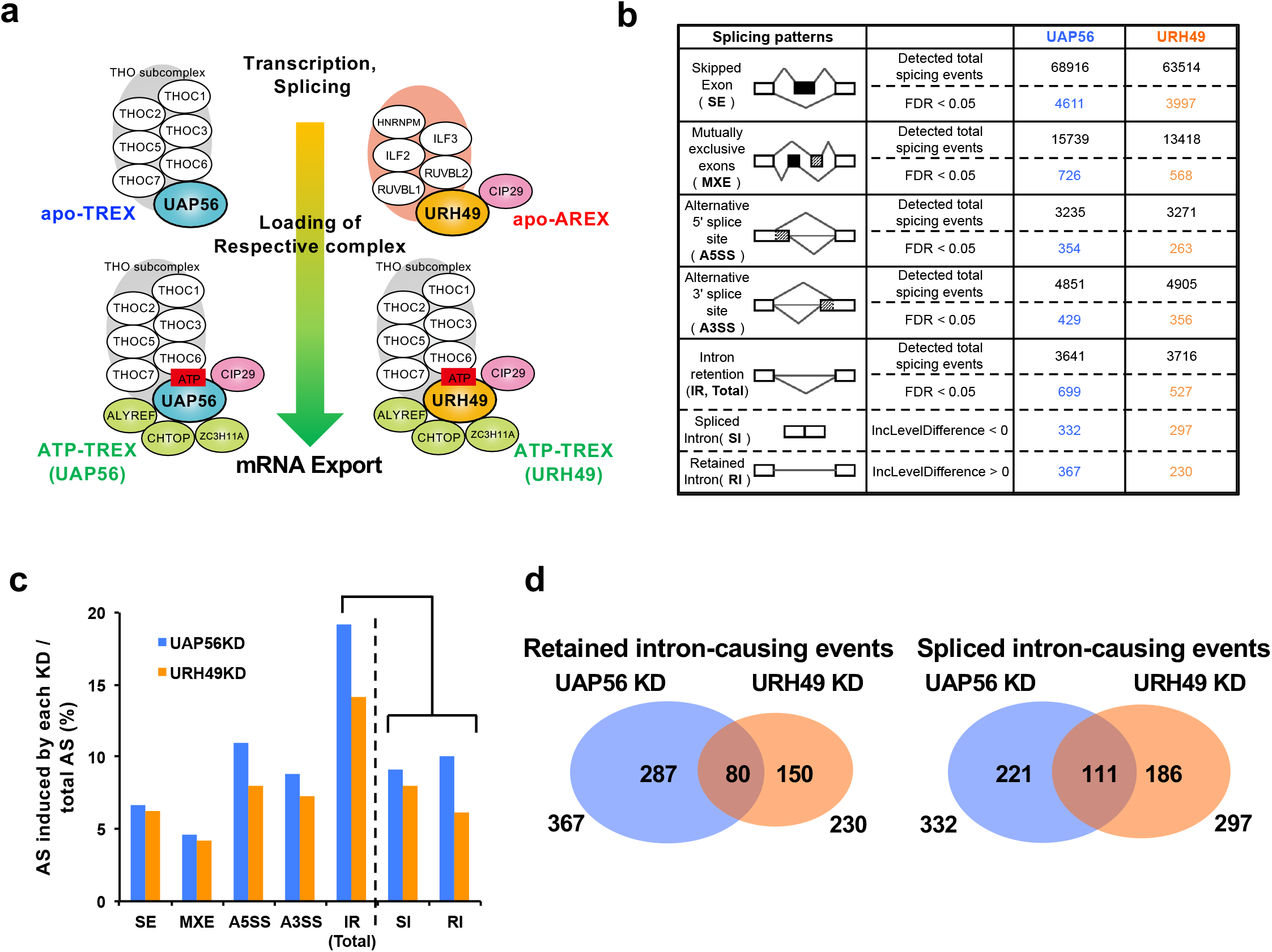
UAP56 and URH49 function in the global splicing regulation. **a** Model of the formation of the apo-TREX and the apo-AREX complexes, and subsequently the remodeling to the ATP-TREX complexes by UAP56 and URH49. **b** Changes in five types of splicing patterns upon depletion of UAP56 or URH49. U2OS cells were transfected with the indicated siRNAs and cultured for 48 hours. Then, total RNA was extracted from the cells for RNA sequencing. Alternative splicing analysis was performed using rMATS. The upper row indicates the number of total splicing events detected in each splicing pattern. The bottom row indicates the number of splicing events that resulted in a statistically significant change (false discovery rate: FDR <0.05) in splicing pattern upon UAP56- (blue) or URH49-knockdown (orange). The category of intron retention (IR) is further divided into two categories, retained intron (RI) and spliced intron (SI). **c** Frequency of splicing changes induced by UAP56- or URH49-knockdown relative to the total splicing events, calculated using the data in **b**, **d** Venn diagram showing the numbers of retained intron and spliced intron events induced by UAP56 or URH49 depletion.

Similar to UAP56, ATP-bound URH49 also forms the ATP-TREX complex (URH49)(5). However, in the absence of ATP, URH49 forms a distinct apo-complex named the apo-<u>A</u>lternative-m<u>R</u>NA-<u>EX</u>port (AREX) complex (Fig.1a)(7,14). Through the formation of their respective apo-complexes and the remodeling into the ATP-TREX complex, the two helicases cooperatively export a subset of mRNAs and selectively export a distinct subset of mRNAs, exhibiting non-redundant roles in mitotic progression and cell proliferation(5,14). Thus, the mRNA export pathways mediated by UAP56 and URH49, along with their respective complex components, contribute to the regulation of gene expression. However, how these helicases and their complexes regulate target transcripts before mRNA export remains unclear.

UAP56 was first identified as a splicing auxiliary factor but studied predominantly as an mRNA exporter later, thus, the contribution of UAP56 to splicing and its relationship with mRNA export remains unclear. During splicing, UAP56 is loaded onto pre-mRNA through interactions with U1 snRNA and U2AF2(15–17), and plays a role in the formation and the remodeling of the spliceosome(18). We previously found that UAP56 and URH49 commonly interact with the core factors of the spliceosome(7). Although several studies have implicated these helicases in splicing(19–22), their broader roles in splicing regulation and the functional relationship between splicing and their respective mRNA export pathways remain poorly understood. We therefore hypothesized that these two helicases, together with their respective complexes, first function during splicing and subsequently promote export of the target mRNAs.

To address this question, we systematically investigated how these helicases and their complexes coordinate mRNA splicing with nuclear export. We found that UAP56 and URH49 exhibit diverse patterns of splicing regulation through their distinct complexes, while coupling splicing to multiple mRNA export pathways. Our findings provide new insights into how closely related RNA helicases contribute to the precise regulation of post-transcriptional gene expression in mammalian cells.

## Methods

### Cell culture

U2OS and HeLa cells were obtained from ATCC (HTB-96, CCL-2), and Flp-In T-REx 293 cells were obtained from Thermo Fisher Scientific (Waltham, MA).

These cells were maintained in Dulbecco’s Modified Eagle’s Medium (Fujifilm Wako, Tokyo, Japan) supplemented with 10% heat-inactivated fetal bovine serum at 37°C.

### Antibodies and antisera

Antibodies and antisera were used as follows: goat anti-lamin B antibody (1:2000 dilution; sc-6216, Santa Cruz Biotechnology, Dallas, TX), mouse anti-GAPDH antibody (1:2000 dilution; 016-25523, Fujifilm Wako), mouse monoclonal anti-β-tubulin antibody (1:2000 dilution; T0198, Sigma-Aldrich Japan), mouse anti-β-actin antibody (1:3000 dilution; A2066, Sigma-Aldrich Japan), mouse monoclonal FLAG M2 antibody (1:3000 dilution; F1804, Sigma-Aldrich Japan, Tokyo, Japan), rabbit anti-PRC1 antibody (1:1000 dilution; 603001, Biolegend, San Diego, CA), rabbit anti-HNRNPM antibody (1:2000 dilution; HPA024344, Sigma-Aldrich Japan), and antisera against THOC1 (1:1000 dilution), THOC5 (1:1000 dilution), ALYREF (1:1000 dilution), CIP29 (1:1000 dilution), NXF1 (1:1000 dilution), DDX19 (1:1000 dilution),UAP56 (1:1000 dilution) and URH49 (1:1000 dilution). Antibodies and antisera used in this study are listed in Supplementary Table.3.

### Plasmids, primers, and siRNAs

To construct the following plasmids, the fragments were obtained by PCR amplification with the addition of restriction enzyme sites or infusion enzyme sites at both ends. pcDNA5-FLAG, pcDNA5-FLAG-UAP56, and pcDNA5-FLAG-URH49 vectors were generated as described previously(5). For the pcDNA5-Halo and pcDNA5-FLAG-Halo, Halo-tags were generated by PCR using the pFN21A plasmid and inserted into pcDNA5 or pcDNA5-FLAG vector, respectively. pcDNA5-Halo-UAP56 and cDNA5-Halo-URH49 vectors were generated by the insertion of their respective open reading frames into pcDNA5-Halo. pcDNA5-FLAG-Halo-UAP56 and cDNA5-FLAG-Halo-URH49 vectors were generated by the insertion of their respective open reading frames into pcDNA5-FLAG-Halo. The construction of the plasmids was confirmed by sequencing. The siRNAs and primers used in this study are listed in Supplementary Table.4 and Supplementary Table.5.

### Transfection of plasmids or siRNA

Transient transfection of siRNA and plasmids was performed using Lipofectamine 2000 (Thermo Fisher Scientific, Waltham, MA) according to the manufacturer’s instructions.

### Isolation of total, cytoplasmic and nuclear RNA

Total RNA was isolated by Sepasol-RNA I super G (Nacalai Tesque, Kyoto, Japan) according to the manufacturer’s instructions. For cytoplasmic RNA preparation, the cells were treated with lysis buffer (20 mM Tris-HCl (pH 8.0), 200 mM NaCl, 1 mM MgCl2, 1% NP-40) on ice for 5 minutes. The cytoplasmic fraction was isolated by brief centrifugation while the nuclear fraction was prepared from the precipitate.

### RNA-sequence (RNA-seq), gene expression and global splicing analysis

U2OS cells were treated with plasmids and/or siRNA transfection and cultured for 48 hours. The preparation of total RNA and cytoplasmic RNA was as described above. RNA-seq libraries were obtained using Truseq Stranded mRNA Library Prep Kit (Illumina, San Diego, CA) or NEBNext Poly(A) mRNA Magnetic Isolation Module NEBNext Ultra II Directional RNA Library Prep Kit (New England Biolabs, Ipswich, MA, USA). RNA-seq was performed with two biological replicates from each group using NextSeq High Output or Illumina NovaSeq X Plus. Sequence reads trimmed for adaptor sequence and low-quality sequence using fastq_quality_trimer and fastq_quality_filter in FASTX-Toolkit (http://hannonlab.cshl.edu/fastx_toolkit/) or TrimGalore version 0.6.5 (http://www.bioinformatics.babraham.ac.uk/projects/trim_galore/), and then mapped to the human hg19 genome using STAR (version2.5.2b)(23) with the parameter -- outFilterType BySJout, --alignEndsType EndToEnd, --outFilterMultimapNmax 20, -- outFilterMismatchNmax 999, --outFilterMismatchNoverReadLmax 0.04, -- alignIntronMin 20, --alignSJoverhangMin 8, --alignSJDBoverhangMin 1 and -- sjdbScore 1. The mapped reads were converted to expression counts for each transcript using RSEM (version 1.2.25)(24) or HTSeq (version 2.0.2) software(25) with a gtf file of GENCODE (version 19). Normalized counts of gene expression for each replicate were obtained by processing EBSeq (version 1.2.25)(26) or DESeq2 software(27).

Alternative splicing was analyzed by rMATS (version 3.2.5)(28). The threshold of false discovery rate (FDR) > 0.05 was considered as nonsignificant and such splicing events were excluded from further analysis. Gene ontology (GO) was performed using the Database for Annotation, Visualization and Integrated Discovery (DAVID: version 6.7)(29). Mapping results were indexed using Samtools (version 1.9)(30) and visualized using Integrative Genomics Viewer (IGV: version 2.3) (http://software.broadinstitute.org/software/igv/home). Splice site and Branch point strength scores were calculated using MaxEntScan(31) and SVM-BP program(32), respectively. For motif analysis of target introns, motifs were calculated using MEME(version 4.9.1)(33). Data analyses were performed with R software(34), and ggplot2(35) was used for data visualization. The raw data from the RNA-seq data have been deposited in the National Center for Biotechnology Information (NCBI) database under accession numbers GSE280484.

### Preparation of PAR-CLIP samples

PAR-CLIP was performed as described previously(36) with some modifications. Briefly, Flp-In T-REx 293 cells expressing Halo-UAP56 or URH49 were labeled with 100 mM 4-SU (Sigma-Aldrich Japan) and then crosslinked on ice by irradiation with UV (365 nm) at 150 mJ/cm^2^. Whole-cell lysate was collected and treated with 5 U/ml RNase T1 (Thermo Fisher Scientific) at 22°C for 15 minutes. Halo-tagged protein was precipitated using HaloLink Resin (Promega Japan, Tokyo, Japan) at 4°C for 5 hours. The resin was then treated with 4 U/μl MNase (NEB Japan, Tokyo, Japan) at 37°C for 5 minutes and subsequently with 0.5 U/ml calf intestinal alkaline phosphatase at 37°C for 10 minutes. Subsequently, a 3’ linker was ligated to the RNA fragments with 0.5 U/ml T4 RNA ligase (Thermo Fisher Scientific) by overnight incubation at 16°C. The RNA fragments were radiolabeled with [γ-^32^P]ATP, and then RNA-UAP56 or URH49 complexes were eluted by cleaving the Halo-tag with TEV Protease (Promega Japan). The eluted samples were loaded onto NuPAGE Novex 4%– 12% Bis-Tris gels (Thermo Fisher Scientific), and the bands corresponding to UAP56 or URH49 were excised. RNA fragments were isolated by electroelution, then treated with proteinase K, and a 5’ linker was ligated on. The 5’ linker-ligated RNAs were loaded onto a 10% TBE-Urea gel (Thermo Fisher Scientific), and the bands ranging from 60 to 100 nt in length were excised. The extracted RNAs were subjected to RT-PCR using P3 primers and the SuperScript III First-Strand Synthesis System (Thermo Fisher Scientific). The resulting cDNAs were amplified by PCR (20–25 cycles) using AccuPrime Pfx SuperMix (Thermo Fisher Scientific) with the primers DSFP5 and DSFP3 as described previously(36). The PCR products ranging from 115 to 175 bp in length were excised, eluted, and sequenced using the Illumina HiSeq 2000 Sequencing System with the primer SSP1. The primers used in this study are listed in Supplementary Table.5.

### Other datasets used

iCLIP data for ALYREF and NXF1 were downloaded from GEO accession GSE113953(37), listed in Supplementary Table.6.

### Processing of PAR-CLIP data, identification and annotation of cluster, and metagene analysis

PAR-CLIP reads were trimmed using Trim Galore (version 0.6.1) and trimmed reads were aligned to human genome assembly (hg19) using STAR algorithm (version 3.5.2b) with the parameter --outFilterMultimapNmax 20. Subsequently, binding sites with T>C transitions were identified using wavClusteR (version 2.24.0)(38), specifically designed for processing PAR-CLIP datasets. For correlation analysis at the gene level, trimmed reads were aligned to hg19 using HTSeq (version 0.11.2)(25). Annotation data for long intergenic non-coding RNA, miscellaneous RNA, piRNA, miRNA, snRNA, snoRNA, mitochondrial tRNA, mitochondrial rRNA, tRNA, rRNA, repetitive elements, and protein-coding transcripts were downloaded from multiple sources described previously(36). The clusters were assigned to the different categories using the above-mentioned data by BEDTools’ IntersectBed (version 2.31.1)(39). Then, the clusters that had been assigned to the category of protein-coding transcript were divided further into CDS, 5’UTR, 3’UTR, and intron. The clusters that had been assigned to multiple categories were assigned a single annotation, as described previously(36). Metagene profiles of PAR-CLIP data across genes were calculated using an in-house script modified from “getMetaGene” function in wavClusteR package. Information regarding gene structure was obtained from TxDb.Hsapiens.UCSC.hg19.knownGene (version 3.2.2)(40) package, and the densities of T>C transition clusters at the 5’ends and 3’ends of exons were summarized as Metagene profiles. The raw data for PAR-CLIP analysis have been deposited in the DNA Data Bank of Japan (DDBJ) database under accession numbers BioProject PRJDB5815.

### Splicing Index analysis

To calculate the splicing index, reads over Exon-Exon junctions and Exon-Intron retentions were counted using IntEREst (version 1.14.0)(41). References of IntEREst were created from GENCODE basic annotation gtf files (version 19). In this study, the splicing index was calculated as:

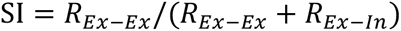

### Quantitative and semi-quantitative reverse transcription-polymerase chain reaction (RT-PCR)

Semi-quantitative RT-PCR was performed as described previously(42). Amplified products were visualized using FAS-IV imaging system (Nippon Genetics, Tokyo, Japan). Quantitative RT-PCR (RT-qPCR) was performed using TB Green Premix Ex Taq II (TakaraBio, Tokyo, Japan) and analyzed by Thermal Cycler Dice real-time system II (TakaraBio). PGK1 was used for standardization. The quantity of each mRNA was calculated based on threshold cycle (Ct) values. The relative amounts of mRNA were evaluated using the values of 2^[Ct (TBP) − Ct (each mRNA)]. Primer sets are listed in Supplementary Table.5. For Splicing efficiency analysis, Pre-mRNA was detected by RT-qPCR using primer extending exon and intron boundary. Spliced mRNA was detected by RT-qPCR using primer extending exon-exon junction. The ratio of pre-mRNA and spliced-mRNA was calculated in each sample. Primer sets are listed in Supplementary Table.5.

### Immunoprecipitation and immunoblotting

Preparation of nuclear extract and immunoprecipitation were performed as described previously(5). Briefly, the nuclear extract was incubated at 20°C for 30 minutes to deplete ATP and centrifuged to recover the supernatant. Then, RNaseA (100 ng/μL) was added to the nuclear extract, and the reaction mixture was incubated at 30°C for 30 minutes. After a brief centrifugation, the clear supernatant was mixed with anti-DYKDDDDK tag antibody beads (Fujifilm Wako) and rotated overnight at 4°C. The beads were extensively washed with PBS containing 0.1% TritonX100, 0.2 mM PMSF, and 0.5 mM DTT to remove nonspecifically bound proteins. The proteins attached to the beads were dissolved in SDS sample buffer (250 mM Tris-HCl, 1% sodium lauryl sulfate), 0.002% bromophenol blue, and 40% glycerol at 37°C for 10 minutes. The eluate was recovered to a new tube and DTT was added to a final concentration of 10 mM, and boiled for 2 minutes.

Samples were separated by SDS-polyacrylamide gel electrophoresis (SDS-PAGE) and blotted to polyvinylidene difluoride membrane (Pall, Ann Arbor, MI). The blot was blocked with PBS containing 0.1% polyoxyethylene sorbitan monolaurate (Tween20) and 5% skimmed milk for 1 hour. The primary antibody was incubated with the blot overnight at 4°C with gentle rotation. The blot was washed extensively with PBS containing 0.1% Tween20. A secondary antibody conjugated with horseradish peroxidase was incubated with the blot for 2 hours with rotation. After extensive washing, the blot was incubated with a chemiluminescence reagent (Millipore, Darmstadt, Germany). Signals were detected using LAS 4000 mini (GE Healthcare Japan, Tokyo, Japan).

### Quantification and statistical analysis

RT-qPCR results were quantified using Thermal Cycler Dice real-time system II (TakaraBio). The statistical significance of two-group and multiple comparisons was determined using R software(34), as indicated in the legends of each Figure. Non-adjusted (two-group comparison) and adjusted (multiple comparison) P-values are presented in each Figure. In box plots, the first and third quartiles are represented by the ends of the box, the median is indicated with a vertical line inside the box, and the minimum and maximum excluding the outliers are represented by the ends of the whiskers. The outliers are indicated by open circles.

## Results

### UAP56 and URH49 function in the global splicing regulation

To investigate the potential role of UAP56 and URH49 in splicing, we examined the transcriptome of U2OS cells depleted UAP56 or URH49. Initially, we validated whether mRNA export is influenced by the suppression of the two helicases. Here, we depleted the respective helicases using siRNAs that had previously been fully rescued from a helicase-deficient mRNA export phenotype by forced expression of FLAG-tagged UAP56 or URH49(7,43). We successfully depleted the two helicases, fractionated the cytoplasmic RNA (Supplementary Fig.1a,b), and evaluated expression changes of transcripts by RNA-sequence (RNA-seq). Consistent with a previous report(14), depletion of either helicase resulted in reduced gene expression in overlapping or different mRNA subsets (Supplementary Fig.1c), indicating that reliable transcriptome analysis was achievable. We also observed increased expression of redundant and specific mRNA subsets upon depleting either helicase (Supplementary Fig.1c).

Next, splicing changes were analyzed using the total RNAs by rMATS(28) (Fig.1b,c, see also Supplementary Table.1). Tens of thousands of splicing events were detected under each knockdown condition and were classified according to the type of splicing alteration. Intron retention category was further divided into two categories, retained intron and spliced intron. Retained intron represents introns with splicing defects and spliced intron represents introns with promoted intron removal. Skipped exon event category can be separated into exon skipping and exon inclusion events. The depletion of either helicase had an impact on all categories of splicing events, suggesting that both helicases globally regulate splicing. To further investigate the relationship between helicase-dependent splicing and mRNA export, we focused primarily on retained intron events, which represent a fundamental class of splicing defects and are the most frequently altered splicing patterns (Fig.1c).

To characterize the splicing events regulated by UAP56 and URH49, we analyzed the genomic features of introns that caused intron retention events. Intron retention events induced by UAP56- and URH49-knockdown were characterized by short length, weak 5’ and 3’ splice sites, and high GC content (Supplementary Fig.1d). The features causing intron retention events by the depletion of either helicase were similar. However, the genomic loci of intron retention and transcripts with splicing changes induced by UAP56 depletion both overlapped with, and in part differed from, those affected by URH49 depletion (Fig.1d, Supplementary Fig.1e). Interestingly, in some introns, the two helicases exhibited opposing effects on splicing (Supplementary Fig.1f). Similar results were observed not only in intron retention category but also exon skipping and exon inclusion categories (Supplementary Fig.1g,h). Gene Ontology enrichment analysis revealed that UAP56- and URH49-depletion partially resulted in splicing changes in gene sets with distinct biological functions (Supplementary Fig.2). These results imply that UAP56 and URH49 cooperatively regulate the splicing of some intron subsets and individually control the splicing of different intron subsets.

### UAP56 and URH49 bind to the pre-mRNAs and spliced-mRNA

To further investigate the regulation of splicing by UAP56 and URH49 *in vivo,* Halo-tagged helicases were used to investigate their distribution on RNA *in vivo* using <u>P</u>hoto <u>A</u>ctivatable-<u>R</u>ibonucleoside-enhanced <u>C</u>ross <u>L</u>inking and <u>I</u>mmuno-<u>P</u>recipitation (PAR-CLIP)(44). We confirmed that each tagged helicase recapitulated the known interactions of the endogenous one (Supplementary Fig.3a) and determined the RNA binding sites of each helicase. The majority of the binding sites of both helicases were categorized protein-coding gene (90 % and 91 % for UAP56 and URH49, respectively, Fig.2a). In the protein-coding gene, UAP56 and URH49 binding clusters were predominantly present in exon (CDS and UTR) and a small fraction (20%) in intron. To elucidate the binding patterns of UAP56 and URH49 to the transcripts, we calculated splicing indexes (Fig.2b). In the UAP56 and URH49 PAR-CLIP read, spliced- and pre-mRNA were found in equal measure. In contrast, the reanalysis of publicly available <u>I</u>ndividual-nucleotide <u>C</u>ross <u>L</u>inking and <u>I</u>mmuno-<u>P</u>recipitation (iCLIP) data for ALYREF and NXF1 revealed that they primarily bind to spliced-mRNA, consistent with previous reports(37). These results raised the possibility that both helicases bind to the pre-mRNAs and remain bound after splicing is completed.

**Fig.2.**
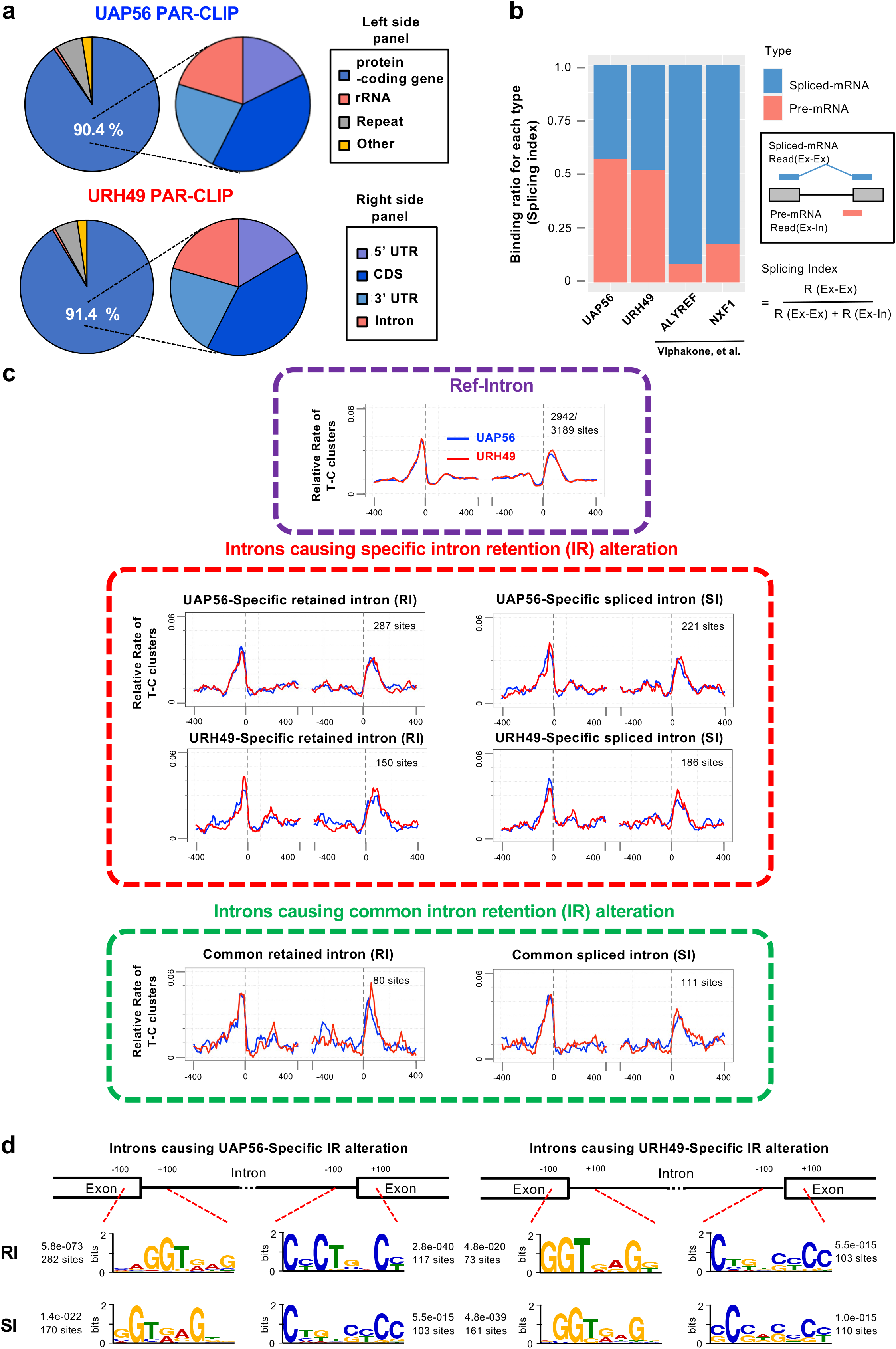
Binding profiles of UAP56 and URH49 onto pre-mRNA and spliced-mRNA. **a** Distribution of binding clusters detected in two replicates of PAR-CLIP reads for UAP56 or URH49. Left side panels: categories of differential RNA species. The “other” category includes long intergenic non-coding RNA, miscellaneous RNA, piRNA, miRNA, snRNA, snoRNA, tRNA, mitochondrial tRNA, and mitochondrial rRNA. Right side panels: regions of protein-coding gene were divided further into coding sequences (CDS), 5’UTR, 3’UTR, and intron as described previously(36). **b** Ratio of the pre-mRNAs to spliced-mRNA in UAP56 or URH49 PAR-CLIP reads. This count was calculated as the ratio of reads crossing internal exon-exon junctions to reads mapped to exon-intron junctions. For comparison, we also reanalyzed the publicly available iCLIP data of ALYREF and NXF1(37). **c** Distribution of the PAR-CLIP reads of UAP56 or URH49 around exon-intron boundaries. The metagene profiles show the occupancy of each binding cluster around the exon-intron boundaries. The left side of each Figure represents the upstream exon-intron boundary, and the right side of each Figure shows the downstream exon-intron boundary. The Ref-Intron indicates the region covering all exons and introns. **d** Representative sequence motif around the exon-intron boundaries of introns with intron retentions caused by depletion of UAP56 or URH49. Consensus sequences were searched using MEME at ±100 bp from the border region of 5’ss and 3’ss of the intron where retained intron or spliced intron was caused by depletion of the two helicases. The Ref-Intron was used as background. The E-value and the number of each motif of the target intron were described.

### Binding profiles of UAP56 and URH49 onto the pre-mRNAs

To establish the transcriptome-wide binding landscape of UAP56 and URH49 to transcripts, we analyzed the distribution of both helicases. The PAR-CLIP reads for UAP56 and URH49 were highly correlated, with only a few transcripts showing different degrees of binding between helicases (Supplementary Fig.3b). Consistently, both helicases showed similar degrees of binding to the UAP56- or URH49-targeted transcripts (Supplementary Fig.3c). These results indicate that the global binding profiles of UAP56 and URH49 were largely similar across the transcriptome. Focusing on the bound position, the peaks of their binding were primarily located around the transcription start site (TSS) and transcription end site (TES) (Supplementary Fig.3d). These binding profiles align with the binding pattern of RNA polymerase II(45), suggesting co-transcriptional recruitments of both helicases. In addition, by focusing on whole exon-intron boundaries, we found that UAP56 and URH49 bound broadly across exon-intron boundaries (Fig.2c), consistent with previous reports on the interactions between UAP56 and U1 snRNA, and U2AF2 *in vitro*(15,17). These results strongly suggested that UAP56 and URH49 play a role in the splicing step.

Next, we examined helicase binding at individual intron and exon–intron boundaries. Although we occasionally observed slightly higher binding of the corresponding helicase near the introns whose retention was altered by helicase depletion (Supplementary Fig.4), these differences were limited. Metagene analyses revealed that UAP56 and URH49 bind to exon–intron boundaries to a similar extent across genome-wide intron regions, introns specifically regulated, or introns cooperatively regulated by either helicase (Fig.2c, Supplementary Table.2). We investigated the relationship between helicase-binding intensity at intron–exon regions surrounding introns with altered splicing upon depletion of either helicase and the degree of splicing changes, but no significant correlation was observed (Supplementary Fig.5). Similarly, in the regions exhibiting changes in the exon skipping, no clear differences were detected in the binding patterns of the two helicases (Supplementary Fig.6).

Finally, we analyzed whether consensus sequences existed for the subset of introns whose intron retention was altered by the depletion of either helicases or their upstream and downstream exons. We found that G- or C-rich motifs in the 5’ or 3’ terminal sequences of the intron, respectively, but their preferences were only slightly different (Fig.2d). No significant motifs were identified within the exons. These results suggest that splicing regulation by UAP56 and URH49 may be determined not by the intrinsic sequence preferences of the helicases, but rather by RNA recognition mechanisms mediated through complex composition, such as the apo-TREX complex or the apo-AREX complex (described in the Discussion section). Taken together, these results suggest that, while modest binding differences can be detected at individual introns, the splicing phenotypes governed by UAP56 and URH49 are not simply dictated by the binding position or intensity, but by additional regulatory layers, including their complex formation and remodeling.

### UAP56 and URH49 coordinate the mRNA export by regulating intron removal

We speculated that the two helicases may coordinate mRNA export through controlling the splicing of target transcripts. To test this possibility, we investigated whether transcripts with intron retention due to the depletion of UAP56 and URH49 altered their cytoplasmic levels (Supplementary Fig.1c). When the transcripts were classified by the subtype of intron removal, transcripts with retained introns were repressed in the cytoplasm by the depletion of the respective helicase (left side panels in Fig. 3a). This indicates that the mRNA export block of target transcripts is determined by whether retained intron events are caused by the depletion of the respective helicase. In contrast, transcripts containing spliced introns were increased in the cytoplasm by the depletion of either one of the two helicases (right panels in Fig.3a). We also showed that depletion of either UAP56 or URH49 promoted mRNA export of some substrates (Supplementary Fig.1c). Our findings indicate that the suppression of splicing due to the depletion of either helicase, as well as its unexpected promotion, is associated with the suppression or promotion of mRNA export. Taken together, these results suggest that UAP56 and URH49 precisely regulate the export of individual transcripts not only by recruiting downstream mRNA export factors shown previously(4,5), but also by newly controlling the removal of introns.

**Fig.3.**
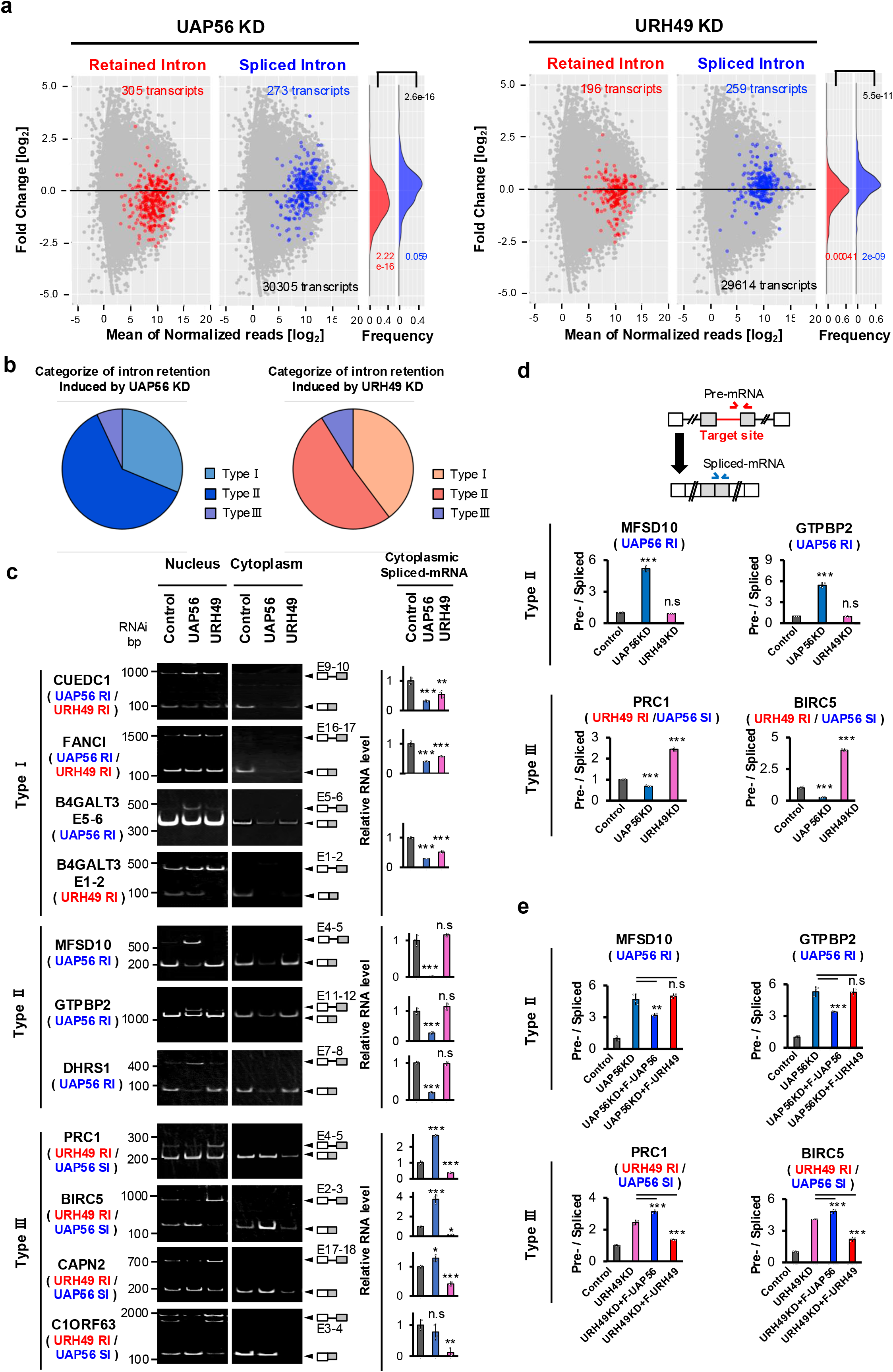
Splicing regulation by UAP56 and URH49 shape multiple mRNA export pathways. **a** Expression changes of cytoplasmic transcripts with retained intron or spliced intron due to depletion of UAP56 and URH49. MA-plot representing the expression change of cytoplasmic transcripts. The log2 fold change of RNA-seq reads upon UAP56 or URH49 depletion is shown on the y axis along with log2 normalized reads on the x axis. Red or blue dots indicate transcripts with retained intron or spliced intron due to depletion of one of the helicases. To focus on the relationship between splicing conditions and their impact on mRNA export, transcripts causing both retained intron and spliced intron upon UAP56 or URH49 depletion were not included in this calculation. The right panels show density plots of the expression changes of transcripts caused by each splicing alteration. Statistical analysis of the gene expression changes for the transcripts with retained introns or spliced introns across the entire gene was performed using t-test. **b** Frequency of each type of splicing alteration to target transcripts upon depletion of either helicase. **c** Depletion of UAP56 or URH49 caused splicing alteration along with the change of their cytoplasmic expression. The left panels show splicing changes detected by RT-PCR using the nucleus and the cytoplasmic RNA. The right panels show mature mRNA expression in the cytoplasm detected by RT-qPCR using cytoplasmic RNA. Values represent the relative expression of the indicated mRNA normalized to PGK1, as mean ±SEM of three independent experiments. Statistical analysis was performed using one-way ANOVA followed by Dunnett’s test. *p < 0.05, **p < 0.01, ***p < 0.001. n.s.: not significant. **d** Either helicase depletion alters splicing efficiency of type II and type III targets. Pre- and spliced mRNA of target transcripts were detected by RT-qPCR. The ratio of pre- to spliced mRNA was calculated. **e** Overexpression of siRNA-resistant FLAG-UAP56 or URH49 specifically rescued splicing defects caused by depletion of UAP56 or URH49. **d, e** Each value is represented as the mean ±SEM of three independent experiments. Statistical analysis was performed using one-way ANOVA followed by Tukey’s test. *p < 0.05, **p < 0.01, ***p < 0.001. n.s.: not significant.

### UAP56 and URH49 regulate splicing to couple multiple mRNA export pathways

Next, we hypothesized that the splicing regulation by UAP56 and URH49 on the transcripts would impact the concertedness and specificity of their mRNA export. To examine this possibility, we classified transcripts with multiple introns into three categories (type I-III) based on the alteration of intron removal upon knockdown of either helicase (Fig.3b, Supplementary Fig.7a,b).

Type I refers to transcripts that require both helicases for efficient splicing and that exhibit a splicing effect in the same direction on intron retention upon depletion of either helicase. This category also included cases in which UAP56 and URH49 regulated distinct introns within the same gene. Type II refers to transcripts that exhibit intron retention upon the depletion of only one helicase. In these transcripts, depletion of one helicase induces intron retention, whereas depletion of another helicase does not lead to detectable changes in splicing. Type III included transcripts for which UAP56 and URH49 exerted opposite effects on splicing. Specifically, depletion of one helicase induced retained intron, whereas depletion of the other promoted intron removal, resulting in antagonistic splicing outcomes (Fig.3b). Each type was also observed when the classification was applied to intron retention events (Supplementary Fig.7c).

We selected several targets and evaluated the correlation between the splicing changes and expression changes of target transcripts in the nucleus and the cytoplasm, in either helicase depletion. In all these situations, intron-containing transcripts were observed in the nuclear and total RNA fractions, but not in cytoplasmic RNA fraction (Fig.3c, Supplementary Fig.8a). In addition, among transcripts with intron retention, spliced-mRNA expression in the cytoplasm was reduced in the case of retained intron transcripts while increased in the case of spliced intron transcripts. These results supported the hypothesis that mRNA export was influenced by the splicing step by the two helicases.

Type I transcripts exhibited reduced cytoplasmic expression upon knockdown of either helicase, indicating that these transcripts had their introns redundantly removed and were exported by either helicase (top category in Fig.3c). Type Ⅱ transcripts showed specific changes in expression only upon the knockdown of either helicase (middle category in Fig.3c). This result suggested that the responsible helicase selectively exported type Ⅱ transcripts. The type III transcripts exhibited opposite changes in splicing and cytoplasmic expression upon knockdown of one helicase (bottom in Fig.3c). These splicing and cytoplasmic expression changes were also confirmed with other cell lines (Supplementary Fig.8b). Consistently, protein expression of PRC1 also changed oppositely by the depletion of respective helicases (Supplementary Fig.8c). These results imply that expression of type III transcripts is tightly and selectively regulated through opposite control of splicing and mRNA export by UAP56 and URH49.

The splicing changes were further confirmed by calculating the ratio of two reverse transcription-quantitative polymerase chain reactions (RT-qPCRs) which amplified pre- and spliced-mRNA (Fig.3d). Crucially, the forced expression of FLAG-helicase resistant to siRNA could rescue these splicing changes induced by the depletion of the corresponding helicase but did not rescue the effect of counterpart helicase depletion (Fig.3e, Supplementary Fig.8d,e). These results indicate that regulation of splicing classified in type I transcripts is associated with cooperative mRNA export, whereas type Ⅱ and Ⅲ transcripts are selectively spliced and exported by UAP56 and URH49.

### Intron-containing transcripts upon UAP56- or URH49-knockdown retained in the nucleus

Premature termination codons (PTCs) emerge from the introns of transcripts with GTPBP2 and PRC1 induced by helicase knockdown. In this context, we examined whether transcripts containing PTCs within retained introns were exported to the cytoplasm and subjected to nonsense-mediated mRNA decay (NMD)(46). In the presence of the NMD inhibitor, cycloheximide (CHX), the SRSF2 transcript variant, a well-known NMD target(47), was efficiently detected in the cytoplasm (Supplementary Fig.9a). In contrast, PTC-containing transcripts induced by the knockdown of UAP56 or URH49 were not detected in the cytoplasm, but were retained in the nucleus (Supplementary Fig.9b). These data indicated that intron-containing transcripts induced by helicase knockdown were retained in the nucleus and blocked mRNA export.

### UAP56 and URH49 regulate transcriptome diversity through the control of alternative splicing

We evaluated the impact of UAP56 and URH49 knockdown on mRNA export efficiency and cytoplasmic expression of alternatively spliced transcripts with skipped exon. The cytoplasmic expression of the transcripts causing skipped exon events was divergently changed by the depletion of UAP56 or URH49 (Fig.4a), indicating a substantial impact of the respective helicase on the skipped exon-containing transcripts. Analogous to intron retained events, skipped exon events were also categorized into three types (type I-III like, Fig.4b). The two helicases regulated various exon skipping or inclusions in a cooperative (type I-like), selective (typeⅡ-like), or antagonistic (type III-like) manner. Inconsistent with the case in retained intron, these alternatively spliced transcripts were efficiently exported to the cytoplasm even in UAP56- or URH49-knockdown condition (Fig.4c). Considering that the two helicases redundantly bind to many transcripts (Supplementary Fig.3b), in the absence of one helicase, the other helicase can function to export alternatively spliced transcripts to the cytoplasm. Taken together, these results indicate that UAP56 and URH49 contribute to the diversity of the transcriptome through regulating the alternative splicing.

**Fig.4.**
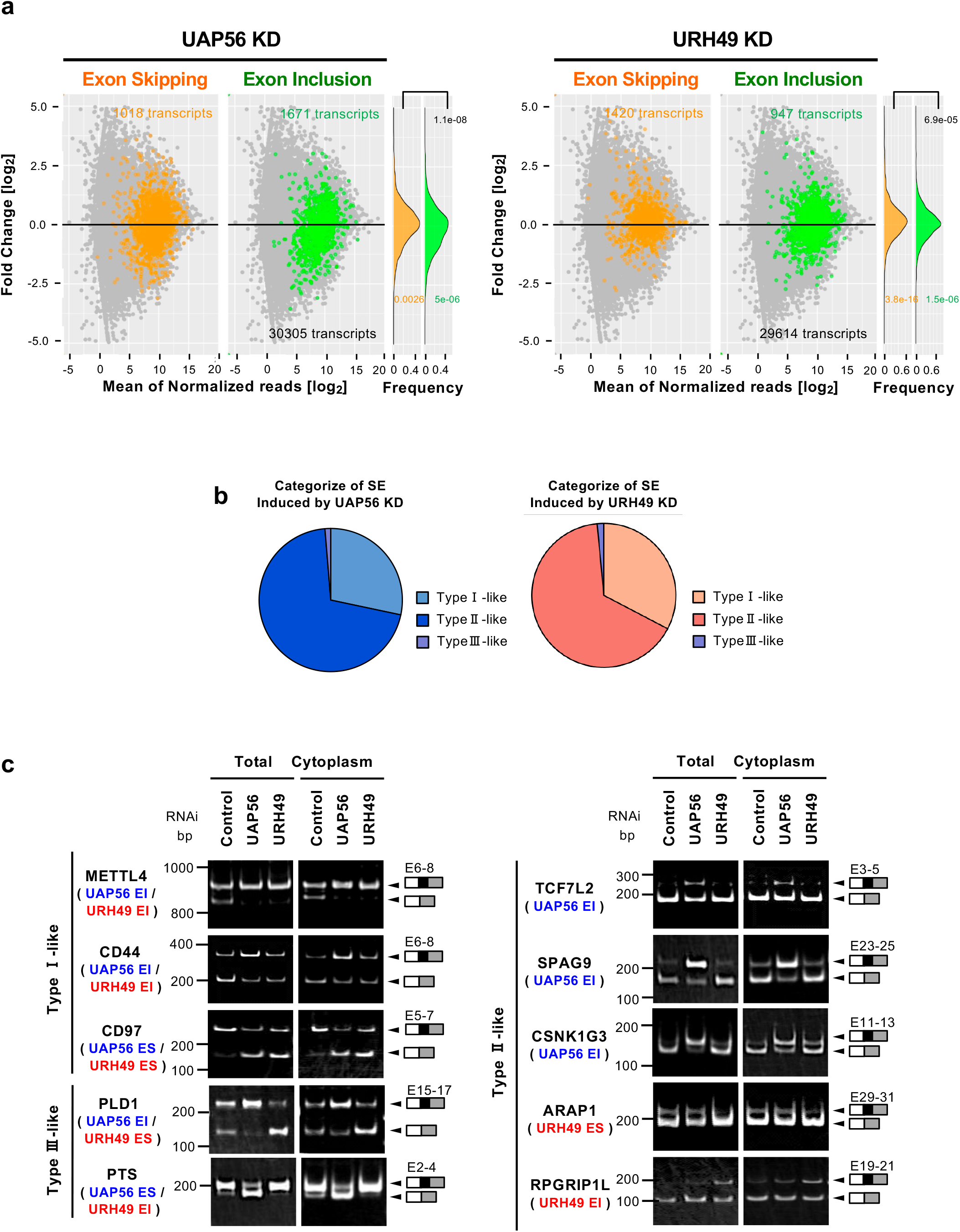
UAP56 and URH49 regulate transcriptome diversity through the control of alternative splicing. **a** Expression changes of cytoplasmic transcripts with exon skipping (ES) and exon inclusion (EI) due to depletion of UAP56 and URH49. MA-plot representing expression changes of cytoplasmic transcripts in the cytoplasm upon depletion of UAP56 or URH49. The log2 fold change in RNA-seq reads upon UAP56 or URH49 depletion is shown on the y axis along with log2 normalized reads on the x axis. Orange or green dots indicate transcripts causing exon skipping or exon inclusion due to depletion of either helicase. To focus on the relationship between splicing conditions and their impact on mRNA export, transcripts causing both exon skipping and exon inclusion upon UAP56 or URH49 depletion were not included in this calculation. The right panels are density plots of the expression changes for transcripts causing each splicing alteration. Statistical analysis of the gene expression changes with exon skipping or exon inclusion across the entire gene was performed using t-test. **b** Frequency of each type of splicing alteration into the target events upon depletion of either helicase. **c** Alternative splicing products induced by UAP56 or URH49 knockdown are exported to the cytoplasm. The panels show splicing changes upon UAP56- or URH49-knockdown detected by RT-PCR using total and cytoplasmic RNA.

### Balance of UAP56 and URH49 regulates splicing for the accurate transcriptome

To further understand the splicing regulation by UAP56 and URH49, we examined the effects of knocking down both helicases simultaneously. Consistent with previous studies, dual knockdown caused more pronounced inhibition of mRNA export than single helicase knockdown (Fig.5a,b)(14). Under these conditions, we analyzed the effects of previously described Type I to Type III splicing substrates (Fig.5c). For Type I introns and Type I-like skipped exons, splicing changes were more pronounced upon dual knockdown than with either helicase alone. For Type II and Type II-like substrates, splicing changes were similar to those observed with individual helicase knockdown.

**Fig.5.**
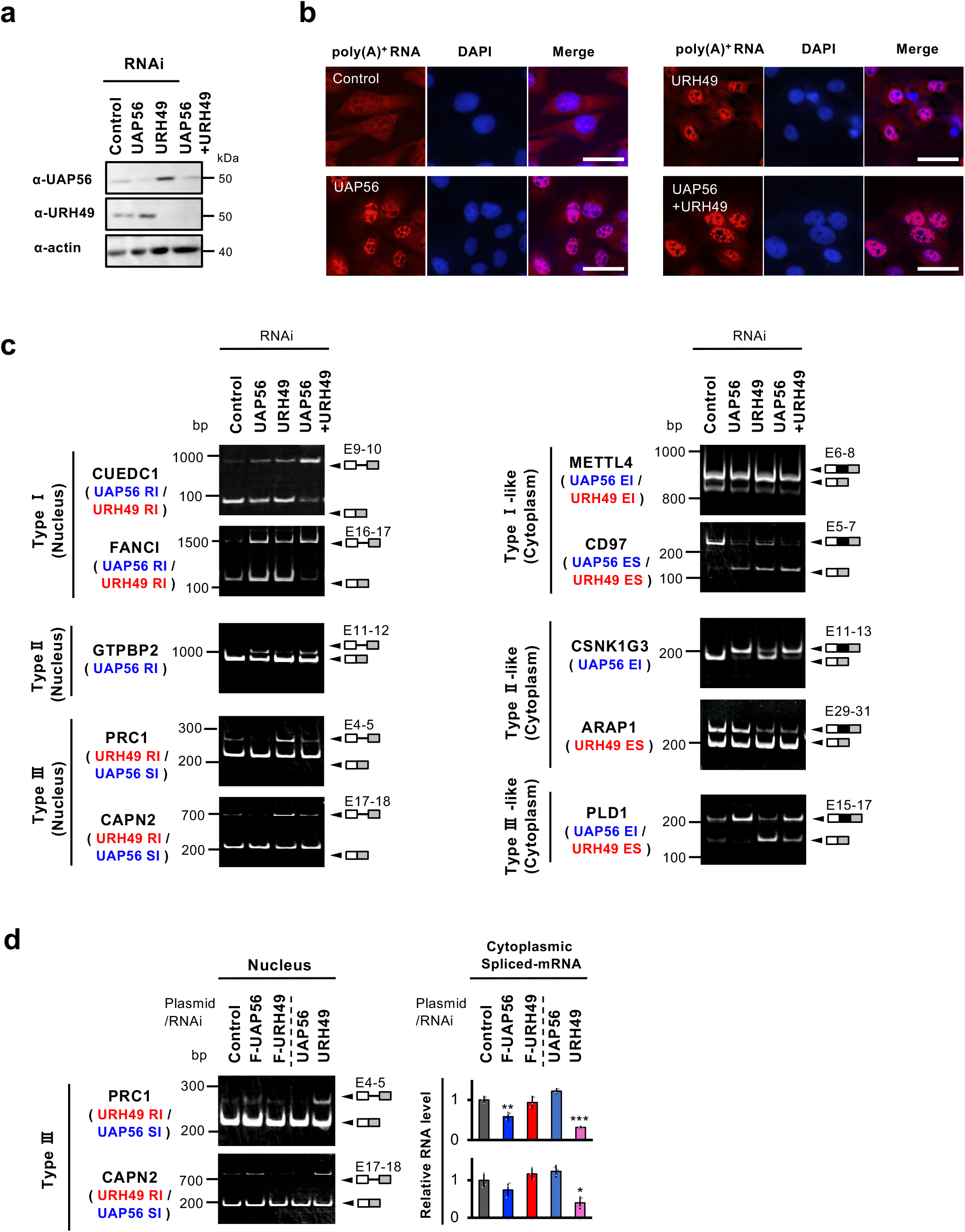
Sophisticated transcriptome regulation by UAP56 and URH49. **a** Knockdown efficiency of individual proteins in U2OS cells was validated using immunoblotting. Actin was used as a loading control. **b** Depletion of UAP56 and URH49 led to nuclear poly(A)^+^ RNA accumulation in U2OS cells. DAPI was used to visualize the nuclei. Scale bar, 40 μm. **c** Splicing changes were induced upon the depletion of UAP56 and URH49. The panels show splicing changes upon UAP56- and URH49-knockdown detected by RT-PCR using nuclear or cytoplasmic RNA. **d** Depletion of UAP56 or URH49 caused splicing alterations along with changes in their cytoplasmic expression. The left panels show splicing changes detected by RT-PCR using nuclear and cytoplasmic RNA. The right panels show the expression of mature mRNA in the cytoplasm as detected by RT-qPCR using cytoplasmic RNA. Values represent the relative expression of the indicated mRNA normalized to PGK1, as mean ±SEM of three independent experiments. Statistical analysis was performed using a one-way ANOVA followed by Dunnett’s test. *p < 0.05, **p < 0.01, ***p < 0.001. n.s.: not significant.

Interestingly, for Type III and Type III-like substrates, dual knockdown abolished the splicing changes observed with single-helicase knockdown. These results support the notion that UAP56 and URH49 cooperatively regulate Type I substrates, selectively regulate Type II substrates, and antagonistically control Type III substrates. To further investigate antagonistic regulation, we examined the effects of forced expression of UAP56 and URH49 (Fig.5d). Forced expression of UAP56, that is, shifting the expression balance between UAP56 and URH49 towards UAP56 dominance, showed effects similar to the mRNA splicing and export effects observed during URH49 knockdown. Taken together, these findings suggest that UAP56 and URH49 coordinate appropriate splicing regulation across Type I to Type III substrates by balancing their expression levels.

### The apo-AREX, and the ATP-TREX complexes are involved in splicing regulation

We analyzed whether the splicing regulation by UAP56 or URH49 is mediated by the formation of apo-TREX complex, apo-AREX complex, and ATP-TREX complex (Fig.1a). We observed that the components of the apo-AREX complex were required for the URH49 targeted intron removal of type I introns (Fig.6a, Supplementary Fig.10a). Furthermore, the depletion of apo-AREX complex components induced splicing defects and down-regulation of the URH49 specific target transcripts, but not the UAP56 specific target transcripts (Fig.6a,b). These results indicated that the apo-AREX complex specifically regulated the splicing of URH49 target transcripts.

**Fig.6.**
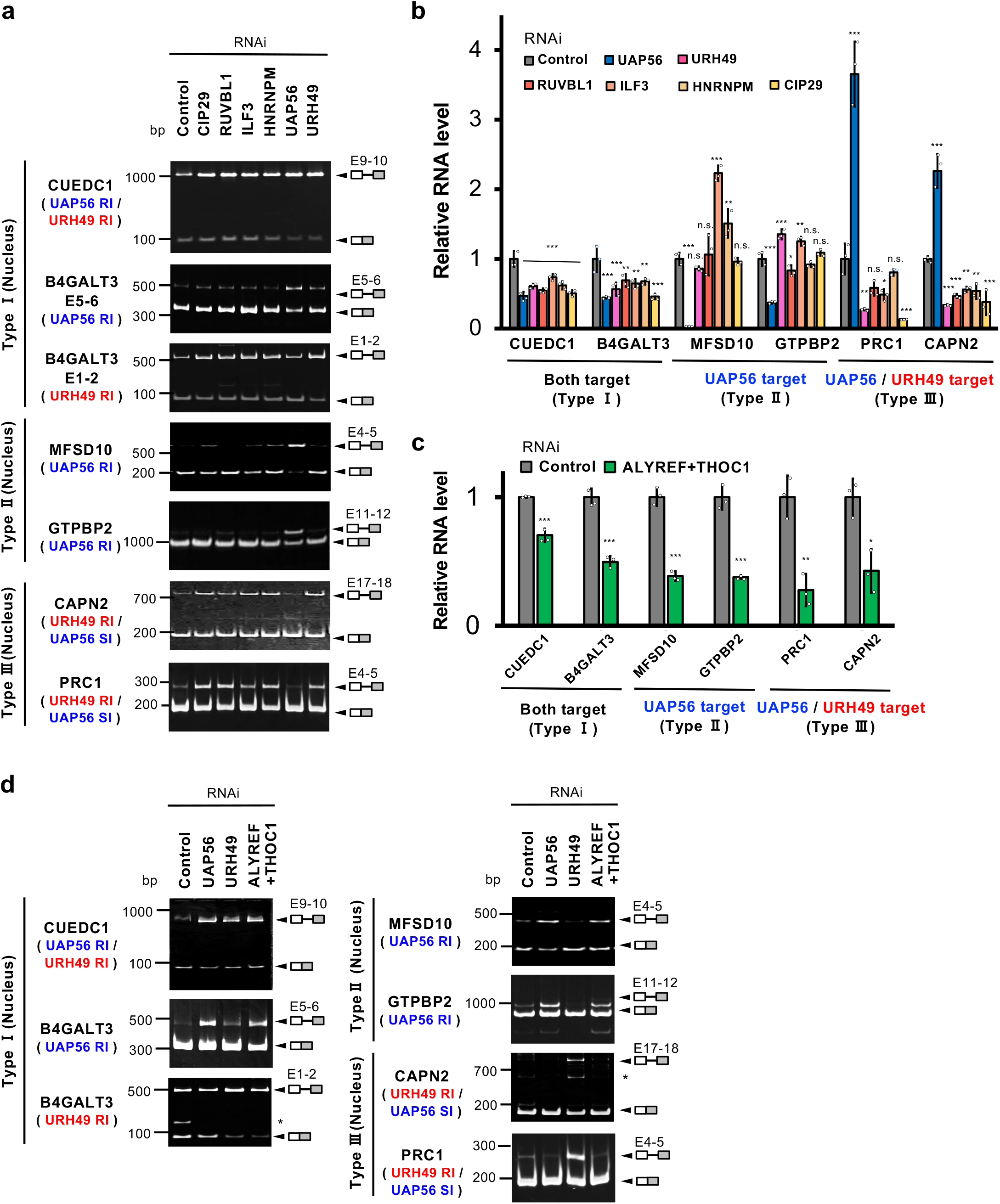
The apo-AREX, and the ATP-TREX complexes regulate splicing. **a** Depletion of the apo-AREX components causes splicing changes similar to the depletion of URH49. The panels depict splicing changes in the nucleus upon the depletion of the apo-AREX components, detected by RT-PCR. **b** Depletion of apo-AREX components downregulates URH49-target mRNAs in the cytoplasm. **c** Depletion of ALYREF and THOC1 downregulates UAP56- and URH49-target mRNAs in the cytoplasm. **b, c** RT-qPCR was performed using the cytoplasmic RNA to calculate the expression level of respective mature mRNA. Values represent the relative expression of the indicated mRNA normalized to PGK1, as the mean ±SEM of three independent experiments. Statistical analysis was performed using one-way ANOVA followed by Dunnett’s test. *p < 0.05, **p < 0.01, ***p < 0.001. n.s.: not significant. **d** Depletion of ALYREF and THOC1 caused splicing changes similar to the depletion of either helicase. The panels depict splicing changes upon the depletion of ALYREF and THOC1 detected by RT-PCR using the nuclear or cytoplasmic RNA.

Since the components of each apo-TREX complex, with the exception of UAP56 or URH49, are also components of the ATP-TREX complex (Fig.1a), the contribution of the apo-TREX complex alone cannot be analyzed directly. Thus, THOC1 and ALYREF were selected as ATP-TREX components in the regulation of splicing by the two helicases, and their contributions were investigated. Depletion of either THOC1 or ALYREF induced the nuclear poly (A)^+^ RNA accumulation (Supplementary Fig.10a,b). The combined depletion of THOC1 and ALYREF provoked a more severe bulk poly (A)^+^ RNA accumulation than either depletion and caused the decreased expression of UAP56 and URH49 target transcripts in the cytoplasm (Fig.6c, Supplementary Fig.10b). With a repressed the ATP-TREX complex, the intron removal categorized as type Ⅰ and type Ⅱ of UAP56 and URH49 target transcripts were inhibited (Fig.6d). The depletion of the ATP-TREX complex components did not affect splicing of the type III of URH49 target intron, thus, their contribution in the splicing of type III intron was not evident. This is probably because the function or formation of the ATP-TREX complex is required for the opposite control of splicing by the two helicases. Similar to the case of intron retention, the depletion of the apo-AREX and the ATP-TREX complexes induced change of alternative splicing regulated by the two helicases, respectively (Supplementary Fig.10c), indicating that the respective complex exhibits a crucial role in the splicing control mediating by UAP56 and URH49. Taken together these results implicate that the apo-AREX complex is involved in URH49-dependent splicing, and the ATP-TREX complex formation is involved in splicing regulation by UAP56 and URH49. We investigated the contribution of apo-complex formation in a later section.

Next, we investigate whether NXF1, a bulk mRNA exporter that interacts with the ATP-TREX complex(5,8–10), contributes to splicing controlled by both helicases. We also examined DDX19, which functioned to release mRNA to the cytoplasm by dissociating mRNA from NXF1 at the cytoplasmic side of the nuclear pore(1). Upon depletion of either factor, an accumulation of bulk poly(A)^+^ RNA in the nucleus was observed (Supplementary Fig.11a,b). Under these conditions, neither NXF1 nor DDX19 were not required for intron removal controlled by UAP56 or URH49, further confirming that the apo-AREX and the ATP-TREX complexes regulate splicing (Supplementary Fig.11c).

### Distinct complex formations are required for the two helicases mediating splicing

DEAD-box RNA helicases have a core region with two domains (N- and C-domains) and terminal regions (Fig.7a)(48). The substitution of their N-domain in UAP56 and URH49 led to changes in their apo-complex formations(7). For example, the UAP56 N-core mutant forms the apo-AREX complex, whereas the URH49 N-core mutant forms the apo-TREX complex. One mutant that forms the apo-TREX complex was particularly noteworthy, namely URH49 C223V, which was generated by mutating the C residue of URH49-C223 to the corresponding V residue of UAP56-V224 within the N-domain.

**Fig.7.**
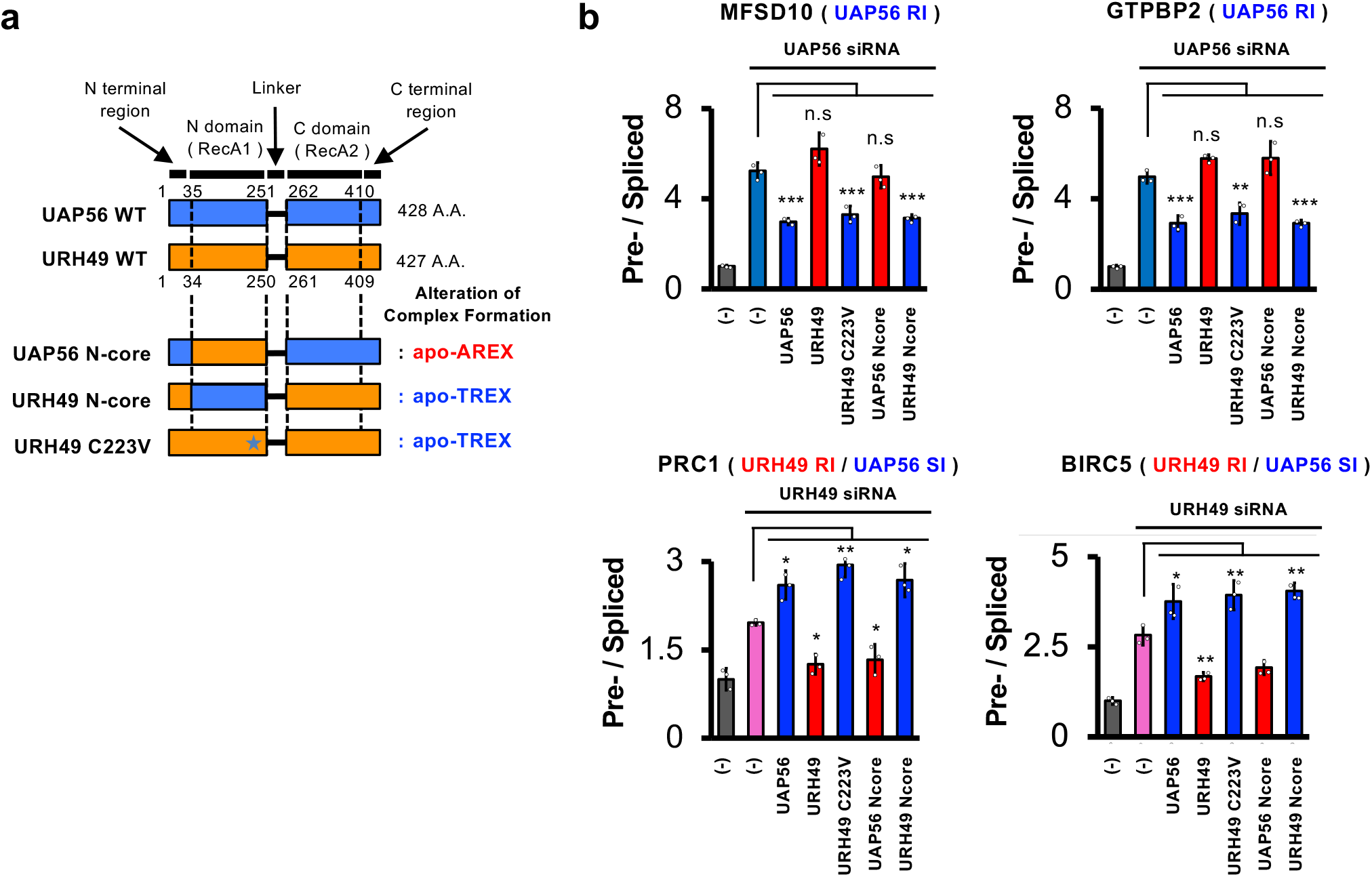
Distinct complex formations are required for the two helicases mediating splicing. **a** Diagram showing the schematic structure of the UAP56 and URH49 with the two RecA regions and the distal regions appropriately numbered. Below the wild type sequences, the chimeric mutants of UAP56 and URH49 are shown along with their apo-complexes, as demonstrated in previous studies(7). **b** Overexpression of chimeric mutants of FLAG-UAP56 or URH49 rescued splicing defects by the respective helicase depletion. Pre- and spliced mRNA of target transcripts were detected by RT-qPCR and the ratio is on the y axis. Each value is represented as the mean ±SEM of three independent experiments. Statistical analysis was performed using one-way ANOVA followed by Tukey’s test. *p < 0.05, **p < 0.01, ***p < 0.001. n.s.: not significant.

To further elucidate the contribution of the distinct apo-complex formation, especially the apo-TREX complex in the splicing regulation by the two helicases, we examined mutants of UAP56 and URH49 for their ability to control the splicing of target transcripts. URH49 C223V and URH49 N-core mutants that form the apo-TREX complex both rescued the splicing defect induced by UAP56 depletion but not the splicing defect induced by URH49 deficiency (Fig.7b). In contrast, the UAP56 N-core mutant that forms the apo-AREX complex specifically rescued the splicing defects induced by URH49 depletion. These data clearly demonstrate that the selective regulation of splicing (type II and type III) by the two helicases is exerted through the formation of their respective apo-complexes.

Integrating insights gained from our study, we propose the following model in which UAP56 and URH49 regulate mRNA splicing and export through their respective complexes. Molecular mechanisms involving complex components, rather than the intrinsic binding properties of helicases alone, contribute to splicing specificity because the formation of apo-TREX, apo-AREX, and ATP-TREX complexes is required for each splicing phenotype. This model is supported by the observation that each helicase binds redundantly to many intron-exon boundaries while controlling splicing in a cooperative, selective, or antagonistic manner. Finally, because the splicing outcomes of transcripts are linked to mRNA export, the export selectivity mediated by UAP56 and URH49 appears to be at least partially coordinated with upstream splicing regulation, thereby contributing to sophisticated gene expression control.

## Discussion

UAP56 and URH49 function in mRNA export along with their respective complexes(5,7,14,43). Both helicases have been shown to play a central role in the remodeling of their respective apo-TREX/AREX complexes into the ATP-TREX complex, and the processes promoted to downstream nuclear export have been elucidated (5,8–13). In addition, recent reports suggest that these two helicases are involved in the splicing regulation of several mRNA subsets(18–22,49). However, it remains unclear whether the individual helicases are involved in general but distinct mRNA splicing throughout the genome. Here, we have revealed a new aspect of their role that the two helicases organize transcriptome accurately by controlling mRNA splicing and export through their respective complexes.

### Closely related helicases have a distinct function in RNA metabolism

The two helicases export mRNAs redundantly and selectively for certain mRNAs(7,14). UAP56 interacts with TREX-2 via the N-terminal region to promote mRNA export(12). On the other hand, URH49 has a weak binding to the TREX-2 complex(12). In addition, UAP56 and URH49 selectively export circular RNAs depending on their lengths(50). URH49 inhibits antiviral activity against alphavirus, while UAP56 does not(51). Thus, these helicases play unique roles in RNA metabolism. We revealed that the two helicases are involved in general splicing, an upstream process of mRNA export, providing a new insight for a better understanding of the distinct functions of the two helicases.

### Precise control of transcriptome via splicing regulation by UAP56 and URH49

The involvement of UAP56 and URH49 in splicing process was provided by their widespread binding to pre-mRNA near exon-intron boundaries. In addition, depletion of either helicase induced splicing changes, suggesting that UAP56 and URH49 primarily control splicing process prior to mRNA export. Although subsets of introns that were either cooperatively or differentially regulated by the two helicases were identified, the overlap of these helicase binding was clearly greater than expected if UAP56 and URH49 acted independently across the genome. This observation indicated that these two helicases acted effectively on many introns in a redundant manner. This interpretation is consistent with their highly similar genome-wide binding profiles, and supports a model in which UAP56 and URH49 share overlapping functions in splicing regulation. A small subset of introns exhibit helicase-specific responses, which cannot be explained by binding profiles alone and are likely determined by additional factors, such as differences in complex composition. These findings are consistent with previous reports indicating that DEAD-box helicases, including UAP56 and URH49, bind to RNA but do not exhibit intrinsic RNA sequence preferences on their own(48).

Interestingly, we observed that both helicases promoted and repressed splicing, suggesting that bidirectional regulation contributes to the precise control of the transcriptome. As a representative example of such context-dependent regulation, we identified antagonistic (type III) splicing controls for PRC1 and BIRC5, which are essential for cell division(52,53). The expression of PRC1 and BIRC5 decreases during the G1 phase and peaks in the G2/M phase(54,55), paralleling the cell-cycle-dependent upregulation of URH49 and CIP29(56,57) whereas UAP56 expression remains relatively constant(57). Consistent with this, changes in the relative expression levels of the two helicases affect the splicing of PRC1 and BIRC5, raising the possibility that transcript expression, such as that of BIRC5 and PRC1, may be fine-tuned through cell cycle-dependent splicing via the balance of UAP56 and URH49. Evaluating how this mechanism functions under physiological conditions may provide new insights into the regulation of cell cycle progression.

Collectively, these findings suggest a novel model in which UAP56 and URH49 regulate splicing based on the balance of their expression levels, thereby contributing to transcriptome regulation. This model offers a functional explanation for the coexistence of two closely related RNA helicases, and suggests that splicing regulation constitutes an upstream layer shaping their downstream roles in mRNA export.

### Role of the apo-TREX, apo-AREX, and the ATP-TREX complex formation in splicing

This study highlights the importance of the complexes formed by UAP56 and URH49 as the mechanistic basis for their involvement in diverse modes of splicing regulation. Furthermore, components of their respective complexes have been identified in purified spliceosomes together with UAP56 and URH49(58–63). These observations support the notion that UAP56 and URH49 do not act on RNA in isolation, but rather function through complexes to engage with the spliceosome and act on pre-mRNAs during splicing. In addition, the UAP56- and URH49-dependent introns identified in this study are GC-rich. This feature is consistent with the previous reports showing that CIP29 and RBM33, which are associated with UAP56 and URH49, preferentially interact with GC-rich RNA sequences(64,65).

These findings raise the possibility that interacting factors within UAP56- and URH49-containing complexes contribute to the selective recognition of target pre-mRNAs. Indeed, in DEAD-box helicase-containing complexes that exhibit RNA sequence-dependent binding, RNA sequence specificity can be conferred by interacting factors rather than by the DEAD-box helicases themselves(48). Determining whether the apo-TREX and apo-AREX complexes exhibit intrinsic RNA sequence preferences would require the isolation and purification of each complex, followed by characterization of their RNA-binding properties. However, their components also participate in other RNA-binding complexes, making it difficult to assess the RNA-binding specificity of individual complexes(7). To address the functional importance of the apo-complex formation from a different perspective, we previously analyzed UAP56 and URH49 mutants with altered abilities to form their respective apo-complexes. Using these mutants, we demonstrated that formation of the apo-TREX and apo-AREX complexes is important for the splicing regulation of their respective target mRNAs. These findings provide evidence that UAP56 and URH49 regulate target mRNA splicing through the formation of the TREX and AREX complexes, respectively, rather than acting solely as individual DEAD-box helicases.

However, it remains unclear how these complexes selectively control the target introns and which cofactors within the complexes are responsible. The reasons why certain introns exhibit such behavior also remain unclear. In particular, the differences in splicing control from type I to type III, or from type I-like to type III-like (Figure 3-6), require additional analysis, including the recruitment of spliceosome factors. To address these concerns, a key future challenge is to elucidate the mechanistic basis by which closely related helicases, through their complex components, acquire distinct specificities for intron selection.

It is also important to consider that complex components can assist helicases by supporting their enzymatic activity. UAP56 regulates spliceosome formation based on its dsRNA helicase and ATPase activities(18), and ATP-TREX components modulate the enzymatic activities of both UAP56 and URH49. The ATP-TREX components, ALYREF, CIP29, and CHTOP, synergistically promote the ATPase and helicase activities of UAP56(10). CIP29 promotes the helicase activity of URH49(66). Taken together with our observations, these findings imply that the apo-TREX, apo-AREX, and ATP-TREX components actively support splicing regulation by facilitating both substrate recognition and enzymatic activity of the two helicases.

### Involvement of UAP56, URH49, and their complex components in cancer

Splicing abnormalities and aberrant gene expression are the hallmarks of tumorigenesis and progression in most cancers(67). URH49 has been found to exhibit abnormally high expression in 18 cancer types, which correlates with these cancer malignancies(68,69), and it has been thought to contribute to cancer-oriented abnormal gene expression through the export of target mRNA. In this study, we revealed that URH49 regulates the splicing of pre-mRNAs from cancer-related genes, such as cell cycle (*p* = 0.02), microtube bundle formation (*p* = 0.033), and positively regulates the canonical Wnt signaling pathway (*p* = 0.049). In addition, our previous studies have shown that URH49 plays an important role in the expression of cell cycle-and cell division–related genes(5,14). Thus, our findings suggest that aberrant expression of URH49 may be associated with the expression of these cancer-related genes through the promotion of their splicing. Up-regulation of UAP56 has also been implicated in tumor progressions in several cancers(19,70,71). We found that UAP56 and URH49 commonly regulate alternative splicing of CD44, which was associated with cancer progression(72). A change in the alternative splicing of HERC4 and SLC37A3 which were thought to be important for cancer progression(73) was also observed by the depletion of either helicase (Supplementary Table.1). These findings suggested that the two helicases might be involved in cancer malignancy through cancer-related gene expression abnormalities including the production of alternatively spliced products. In addition, the abnormal expression of the apo-TREX, the apo-AREX, and the ATP-TREX components has also been reported in various cancers(4), implicating that respective complex is predicted to be involved in transcriptome abnormalities in cancer through splicing. Further insight into UAP56 and URH49 and their complexes will advance our understanding of the transcriptome changes during various cancer progressions.

### Conserved role of the splicing regulation during evolution

During evolution, many RNA-binding proteins have functionally diversified to shape diverse transcriptomes contributing to the complexity of living organisms.

Sub2, a homolog of UAP56 in yeast, also plays a role in splicing(74–76). In *Arabidopsis thaliana*, THOC3, a homolog of the THO component, is physically associated with spliceosomes(77). Moreover, THOC1, THOC2, and THOC3 are required for several alternative splicing events(77–79). These findings indicated that the association of UAP56 and the apo- and ATP-TREX components in splicing is conserved within various species of animals and plants. Furthermore, the ortholog of URH49 was diversified from an ancestor of UAP56 during the evolution of vertebrates and invertebrates(7). Thus, it is likely that UAP56 and URH49, along with their respective complexes, have contributed to the acquisition of splicing regulatory networks for the precise control of transcriptome in a variety of organisms, not limited to humans.

## Data availability

RNA-seq data have been deposited in the NCBI (GSE280484) and PAR-CLIP data have been deposited in the DDBJ database (BioProject PRJDB5815).

## Supplementary Data statement

Supplementary data is available at NAR online.

## Acknowledgments

We thank Dr. Takefumi Kondo for technical support in RNA-seq analysis, NXF1 anti-sera for kindly gift from Dr. Ichiro Taniguchi, DDX19 anti-sera for kindly gift from Dr. Robin Reed, Dr. Kei Iida and Dr. Tomohiro Yamazaki for helpful suggestions and encouragement, and our lab members for their constructive discussions. Some computations were performed using the NIG supercomputer at National Institute of Genetics, Mishima, Japan. English proofreading was performed by Editage.

## Author contributions statement

K.F. and S.M. conceived and designed this study; K.F. performed the experiments and analyses including bioinformatics analyses, organized the data and drafted the manuscript; K.H, Y.I, H.Y, and F.H, assisted with biological experiments; S.S. and H.M. performed bioinformatics analyses; M.Y. and K.Y. performed PAR-CLIP sample preparation; K.F., N.F, A.M, and S.M. wrote the paper. All authors reviewed the final manuscript.

## Funding

This work was supported in part by Grants-in-Aid from JSPS KAKENHI [26292053, 221S0002, 17K19232, 19K22280, 19H02884, 21K19078, 22H02264, 23K23531, 16H06279, 22H04925, 25H00433 and 25K24726 to S.M, 19K15807 to K.F]. This work was also supported in part by Grants-in-Aid from Agricultural Technology and Innovation Research Institute, Kindai University, Antiaging Center, Kindai University and The Mitsubishi Foundation to S.M and The Sasakawa Scientific Research Grant from The Japan Science Society to K.F.

## Conflict of interest disclosure

The authors declare no competing interests.

## Notes

### Competing Interest Statement

The authors have declared no competing interest.

