## Supplementary Figures and Tables for "Two closely related RNA helicases, UAP56 and URH49, couple mRNA splicing and export pathways with their distinct complexes"

### SFigure.1

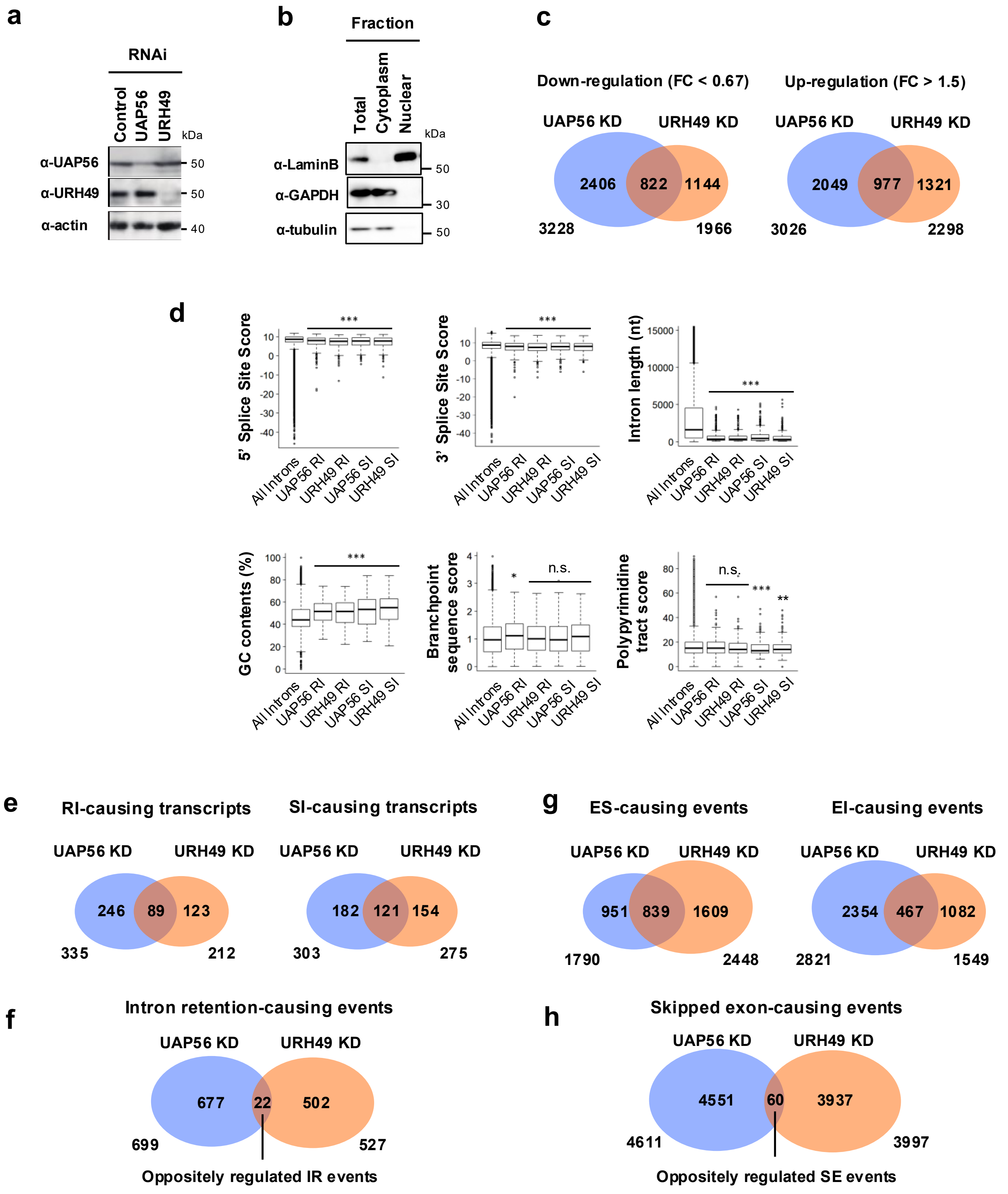
a
RNAi
Control
UAP56
URH49
kDa
α-UAP56
α-URH49
α-actin
50
50
40
b
Fraction
Total
Cytoplasm
Nuclear
kDa
α-LaminB
α-GAPDH
α-tubulin
50
30
50
c
Down-regulation (FC < 0.67)
UAP56 KD
URH49 KD
2406
822
1144
3228
1966
Up-regulation (FC > 1.5)
UAP56 KD
URH49 KD
2049
977
1321
3026
2298
d
5' Splice Site Score
3' Splice Site Score
Intron length (nt)
GC contents (%)
Branchpoint sequence score
Polypyrimidine tract score
All Introns
UAP56 RI
URH49 RI
UAP56 SI
URH49 SI
***
*
**
n.s.
e
RI-causing transcripts
UAP56 KD
URH49 KD
246
89
123
335
212
SI-causing transcripts
UAP56 KD
URH49 KD
182
121
154
303
275
g
ES-causing events
UAP56 KD
URH49 KD
951
839
1609
1790
2448
EI-causing events
UAP56 KD
URH49 KD
2354
467
1082
2821
1549
f
Intron retention-causing events
UAP56 KD
URH49 KD
677
22
502
699
527
Oppositely regulated IR events
h
Skipped exon-causing events
UAP56 KD
URH49 KD
4551
60
3937
4611
3997
Oppositely regulated SE events

**Supplementary Figures and Tables**

**Supplementary Fig.1. Feature of splicing changes induced by the depletion of UAP56 and URH49 (Related to Fig.1)**

**a** Knockdown efficiency of individual proteins in U2OS cells was validated by immunoblotting. Actin was used as a loading control. **b** Fractionations of the nuclear and the cytoplasmic RNA were confirmed by immunoblotting. GAPDH and tubulin were used as cytoplasmic markers, and Lamin B was used as a nuclear marker.

**c** Venn diagram illustrating the number of transcripts with altered expression in the cytoplasm upon knockdown of UAP56 or URH49. Respective gene expression was analyzed by RSEM and EBSeq. **d** Bioinformatic analysis of introns with retained or spliced introns upon the depletion of UAP56 or URH49. All introns of human genes were analyzed for comparison. Boxes represent the median (center line) and upper and lower quartile ranges. Whiskers indicate minimum and maximum values. Statistical analysis was performed using the Kruskal-Wallis followed by Steel test. *$p < 0.05$, **$p < 0.01$, ***$p < 0.001$, n.s.: not significant. **e** Venn diagram represents the numbers of transcripts with retained intron (RI) or spliced intron (SI) under the UAP56- or URH49-depleted condition. **f** Venn diagram showing the numbers of intron retention (IR) events and oppositely regulated IR events induced by UAP56 or URH49 depletion. **g** Venn diagram showing the numbers of exon skipping (ES) and exon inclusion (EI) events caused by UAP56- or URH49-depletion. **h** Venn diagram showing the numbers of skipped exon (SE) events and oppositely regulated SE events induced by UAP56 or URH49 depletion.

### SFigure.2

## RI

##### UAP56 KD

| | Term | Count | P-Value |
|---|---|---|---|
| * | DNA repair | 17 | 3.6E-06 |
| * | DNA recombination | 9 | 0.000095 |
| * | regulation of signal transduction by p53 class mediator | 9 | 0.0014 |
| | RNA splicing | 9 | 0.0085 |
| * | cell volume homeostasis | 3 | 0.012 |
| | RNA export from nucleus | 5 | 0.015 |
| * | IRE1-mediated unfolded protein response | 5 | 0.019 |
| * | mitotic G2 DNA damage checkpoint | 3 | 0.027 |
| * | negative regulation of telomerase activity | 3 | 0.027 |
| * | tRNA aminoacylation for protein translation | 4 | 0.032 |
| * | Golgi organization | 5 | 0.04 |
| | mRNA splicing, via spliceosome | 9 | 0.04 |
| * | RNA secondary structure unwinding | 4 | 0.04 |
| | transcription, DNA-templated | 45 | 0.042 |
| | cell division | 12 | 0.043 |

##### URH49 KD

| | Term | Count | P-Value |
|---|---|---|---|
| * | RNA export from nucleus | 6 | 0.00041 |
| * | protein sumoylation | 7 | 0.0023 |
| * | mRNA 3'-end processing | 5 | 0.0025 |
| | RNA splicing | 8 | 0.003 |
| | mRNA splicing, via spliceosome | 9 | 0.0041 |
| | mRNA export from nucleus | 6 | 0.0058 |
| * | termination of RNA polymerase II transcription | 5 | 0.0062 |
| * | mRNA processing | 7 | 0.017 |
| | cell division | 10 | 0.02 |
| * | microtubule bundle formation | 3 | 0.033 |
| | positive regulation of transcription, DNA-templated | 12 | 0.035 |
| * | viral transcription | 5 | 0.04 |
| * | adenosine to inosine editing | 2 | 0.045 |
| * | negative regulation of ubiquitin-protein ligase activity involved in mitotic cell cycle | 4 | 0.048 |
| * | positive regulation of canonical Wnt signaling pathway | 5 | 0.049 |

## SI

| | Term | Count | P-Value |
|---|---|---|---|
| | mRNA splicing, via spliceosome | 13 | 0.00023 |
| | regulation of translational initiation | 6 | 0.00025 |
| | translational initiation | 10 | 0.00035 |
| * | protein phosphorylation | 18 | 0.001 |
| * | IRE1-mediated unfolded protein response | 6 | 0.0024 |
| * | positive regulation of canonical Wnt signaling pathway | 8 | 0.0031 |
| * | proteasome-mediated ubiquitin-dependent protein catabolic process | 10 | 0.0053 |
| * | exonucleolytic trimming to generate mature 3'-end of 5.8S rRNA | 3 | 0.0066 |
| * | nuclear mRNA surveillance | 3 | 0.0084 |
| | intraciliary retrograde transport | 3 | 0.01 |
| * | regulation of mitophagy | 4 | 0.015 |
| | RNA splicing | 8 | 0.017 |
| * | termination of RNA polymerase II transcription | 5 | 0.019 |
| * | rRNA processing | 9 | 0.021 |
| * | cilium morphogenesis | 7 | 0.022 |

| | Term | Count | P-Value |
|---|---|---|---|
| | mRNA splicing, via spliceosome | 14 | 0.000021 |
| * | histone H3 deacetylation | 4 | 0.0032 |
| * | RNA secondary structure unwinding | 5 | 0.0036 |
| * | nuclear-transcribed mRNA poly(A) tail shortening | 4 | 0.0082 |
| * | protein deacetylation | 3 | 0.0086 |
| | intraciliary retrograde transport | 3 | 0.0086 |
| * | protein homooligomerization | 8 | 0.014 |
| | translational initiation | 7 | 0.014 |
| * | mRNA export from nucleus | 6 | 0.015 |
| * | phospholipid biosynthetic process | 4 | 0.02 |
| * | covalent chromatin modification | 6 | 0.024 |
| * | histone H3 acetylation | 4 | 0.024 |
| * | fatty acid beta-oxidation using acyl-CoA dehydrogenase | 3 | 0.027 |
| * | positive regulation of chaperone-mediated protein complex assembly | 2 | 0.029 |
| * | gene expression | 4 | 0.032 |

## EI

| | Term | Count | P-Value |
|---|---|---|---|
| | cell division | 52 | 7.10E-08 |
| | cilium morphogenesis | 27 | 7.40E-07 |
| | cilium assembly | 23 | 1.80E-05 |
| | intracellular protein transport | 34 | 3.30E-05 |
| | mitotic nuclear division | 34 | 9.00E-05 |
| * | nucleotide-binding oligomerization domain containing signaling pathway | 9 | 1.40E-04 |
| * | regulation of mitophagy | 10 | 2.10E-04 |
| | G2/M transition of mitotic cell cycle | 22 | 2.40E-04 |
| | sister chromatid cohesion | 18 | 3.80E-04 |
| * | protein localization to cilium | 7 | 7.50E-04 |
| * | protein transport | 44 | 8.10E-04 |
| | proteasome-mediated ubiquitin-dependent protein catabolic process | 27 | 8.60E-04 |
| * | vesicle-mediated transport | 22 | 1.00E-03 |
| | protein polyubiquitination | 25 | 1.00E-03 |
| * | double-strand break repair | 13 | 1.10E-03 |

| | Term | Count | P-Value |
|---|---|---|---|
| | G2/M transition of mitotic cell cycle | 35 | 1.90E-09 |
| * | RNA splicing | 36 | 9.80E-08 |
| | protein transport | 63 | 2.40E-07 |
| | cilium assembly | 28 | 1.40E-06 |
| * | mRNA processing | 35 | 1.90E-06 |
| | ubiquitin-dependent protein catabolic process | 34 | 7.60E-06 |
| | cell division | 52 | 2.30E-05 |
| | cilium morphogenesis | 27 | 2.70E-05 |
| | sister chromatid cohesion | 22 | 5.80E-05 |
| | protein ubiquitination involved in ubiquitin-dependent protein catabolic process | 28 | 8.10E-05 |
| * | centriole replication | 8 | 1.30E-04 |
| * | telomere maintenance via recombination | 11 | 1.30E-04 |
| * | cell-cell adhesion | 40 | 2.60E-04 |
| | proteasome-mediated ubiquitin-dependent protein catabolic process | 32 | 3.90E-04 |
| | protein autoubiquitination | 13 | 4.70E-04 |

## ES

| | Term | Count | P-Value |
|---|---|---|---|
| * | mRNA splicing, via spliceosome | 54 | 4.90E-10 |
| * | RNA splicing | 43 | 4.90E-09 |
| | DNA repair | 53 | 1.20E-08 |
| | mRNA processing | 43 | 5.10E-08 |
| | cell division | 67 | 1.10E-07 |
| | G2/M transition of mitotic cell cycle | 35 | 2.30E-07 |
| * | DNA replication | 36 | 1.70E-06 |
| | mitotic nuclear division | 49 | 2.70E-06 |
| | cell-cell adhesion | 52 | 3.10E-06 |
| * | cell cycle | 43 | 1.10E-05 |
| | protein autoubiquitination | 17 | 1.20E-05 |
| | sister chromatid cohesion | 26 | 1.30E-05 |
| * | mRNA export from nucleus | 25 | 2.40E-05 |
| * | termination of RNA polymerase II transcription | 19 | 2.60E-05 |
| | RNA processing | 24 | 4.30E-05 |

| | Term | Count | P-Value |
|---|---|---|---|
| | DNA repair | 32 | 6.70E-06 |
| | protein polyubiquitination | 26 | 3.10E-05 |
| * | DNA damage response, signal transduction by p53 class mediator resulting in cell cycle arrest | 13 | 1.30E-04 |
| | cell-cell adhesion | 31 | 2.50E-04 |
| * | regulation of epithelial to mesenchymal transition | 6 | 3.00E-04 |
| | G2/M transition of mitotic cell cycle | 19 | 5.80E-04 |
| * | establishment of mitotic spindle localization | 4 | 6.50E-04 |
| | regulation of mRNA processing | 5 | 9.50E-04 |
| * | positive regulation of basement membrane assembly involved in embryonic body morphogenesis | 4 | 1.60E-03 |
| * | protein phosphorylation | 42 | 1.60E-03 |
| | proteasome-mediated ubiquitin-dependent protein catabolic process | 23 | 2.10E-03 |
| | cell division | 34 | 2.10E-03 |
| * | cytoskeletal anchoring at plasma membrane | 5 | 2.30E-03 |
| | cellular response to DNA damage stimulus | 23 | 2.80E-03 |
| | mitotic nuclear division | 26 | 2.90E-03 |

**Supplementary Fig.2. Gene ontologies of splicing changes induced by the depletion of UAP56 and URH49 (Related to Fig.1)**

Transcripts with retained intron, spliced intron, exon skipping, and exon inclusion caused by UAP56 or URH49 depletion were enriched for several gene ontology terms categorized in “Biological Process”. Each term is listed in order of increasing p-value. Count indicates the number of genes caused retained intron, spliced intron, exon skipping, and exon inclusion by UAP56 or URH49 knockdown.

### SFigure.3

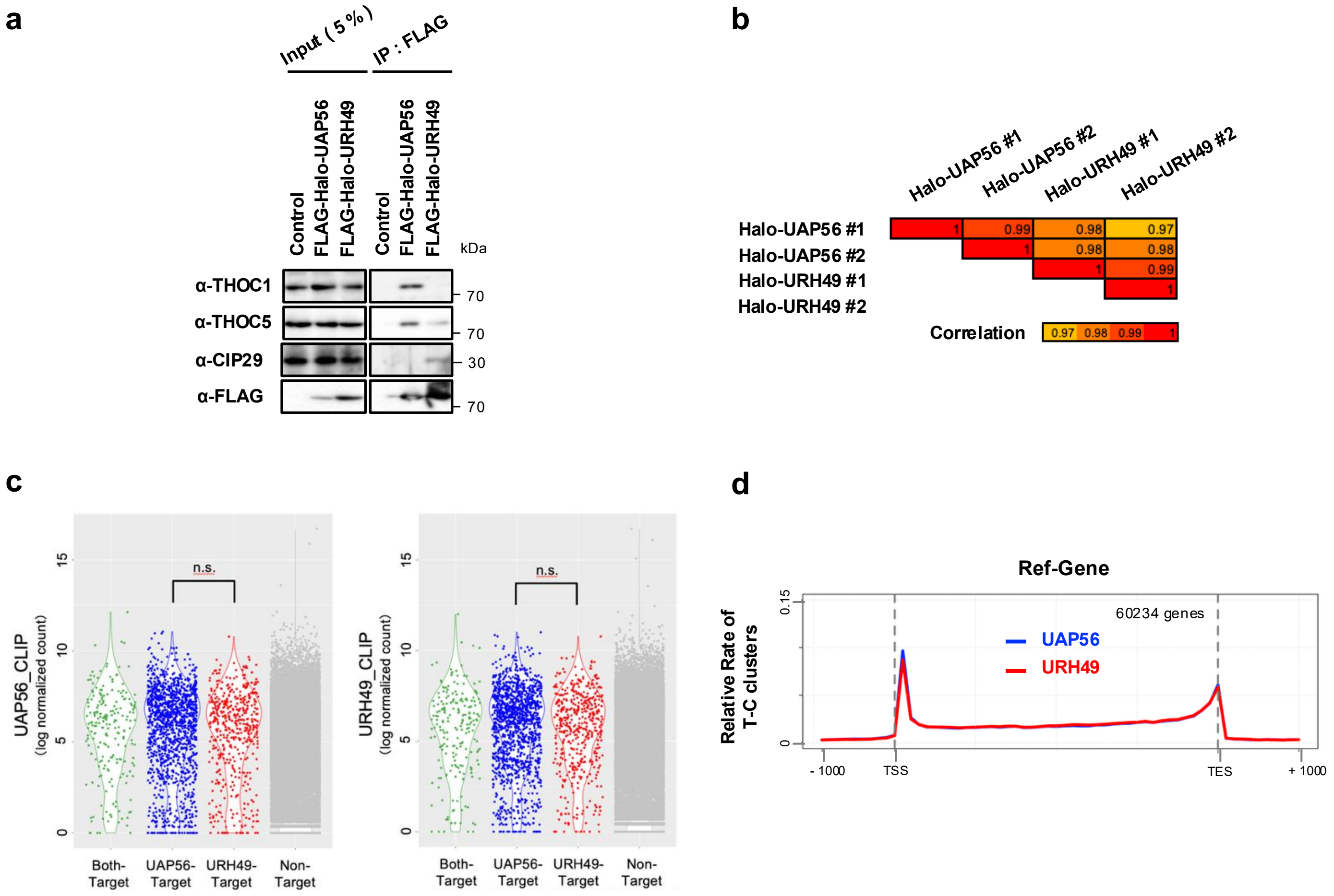
a
Input ( 5 % )
IP : FLAG
Control
FLAG-Halo-UAP56
FLAG-Halo-URH49
Control
FLAG-Halo-UAP56
FLAG-Halo-URH49
kDa
α-THOC1
α-THOC5
α-CIP29
α-FLAG
70
70
30
70
b
Halo-UAP56 #1
Halo-UAP56 #2
Halo-URH49 #1
Halo-URH49 #2
Halo-UAP56 #1
Halo-UAP56 #2
Halo-URH49 #1
Halo-URH49 #2
1
0.99
0.98
0.97
1
0.98
0.98
1
0.99
1
Correlation
0.97
0.98
0.99
1
c
UAP56_CLIP
(log normalized count)
URH49_CLIP
(log normalized count)
n.s.
n.s.
0
5
10
15
Both-Target
UAP56-Target
URH49-Target
Non-Target
d
Ref-Gene
60234 genes
UAP56
URH49
Relative Rate of T-C clusters
0
0.15
- 1000
TSS
TES
+ 1000

**Supplementary Fig.3. Feature of the RNA binding pattern of UAP56 and URH49 (Related to Fig.2)**

**a** Halo- and FLAG-tagged UAP56 or URH49 associated with the apo-TREX and AREX components. Immunoprecipitation was performed using anti-DYKDDDDK tag antibody beads. Precipitated samples were separated and detected by immunoblotting with the indicated antibodies. **b** Correlation of UAP56 and URH49 PAR-CLIP reads. We analyzed the gene expression level of the respective PAR-CLIP sample analyzed by HTSeq. The Pearson correlation coefficients were represented as a heatmap. **c** Violin plots showing the intensities of each CLIP signal for UAP56- or URH49-target transcripts. The signal intensity was calculated using the HTSeq count and DESeq2 software. Statistical analysis was performed using Student's t-test, and pairwise comparisons between levels of each group were calculated using Bonferroni corrections for multiple testing. n.s.: not significant. From the RNA-seq reads upon UAP56 or URH49 depletion, the expression levels of each mRNA were analyzed using RSEM and EBSeq. mRNAs with expression levels specifically reduced by more than twofold due to UAP56 and URH49 depletion were defined as UAP56- or URH49-target mRNAs. **d** Distribution of the binding clusters of UAP56 or URH49 across the entire transcriptome, detected and analyzed by waveClusteR.

### SFigure.4

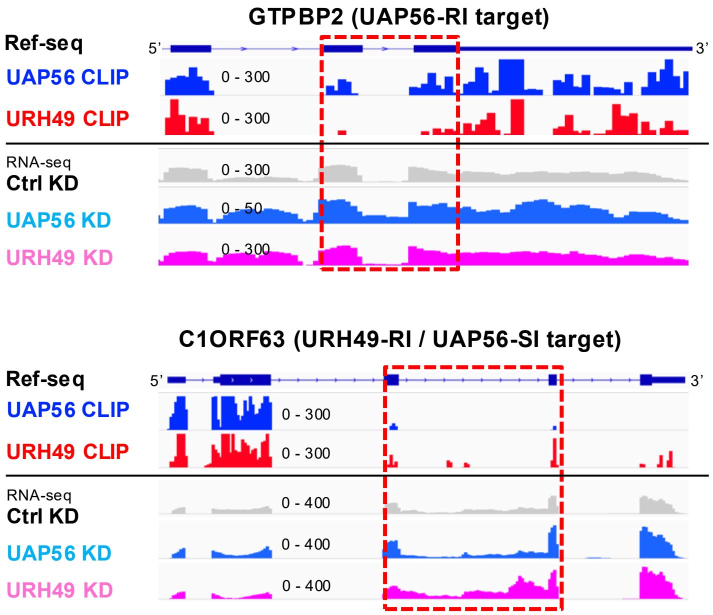
GTPBP2 (UAP56-RI target)
Ref-seq
5'
3'
UAP56 CLIP
0 - 300
URH49 CLIP
0 - 300
RNA-seq
Ctrl KD
0 - 300
UAP56 KD
0 - 50
URH49 KD
0 - 300
C1ORF63 (URH49-RI / UAP56-SI target)
Ref-seq
5'
3'
UAP56 CLIP
0 - 300
URH49 CLIP
0 - 300
RNA-seq
Ctrl KD
0 - 400
UAP56 KD
0 - 400
URH49 KD
0 - 400

**Supplementary Fig.4. The RNA binding pattern of UAP56 and URH49 and their involvement in splicing (Related to Fig.2)**

Representative differences in the binding of UAP56 and URH49 and the status of intron removal of GTPBP2 or C1ORF63 transcripts in which the depletion of UAP56 and URH49 caused splicing changes. Dashed boxes indicate the exon-intron boundaries of particular introns with differences in splicing regulation. The top line indicates exon and intron structures. The second and third lines represent normalized PAR-CLIP reads of UAP56 or URH49 and the lower three lines represent normalized RNA-seq reads from control UAP56 or URH49 knockdown cells.

### SFigure.5

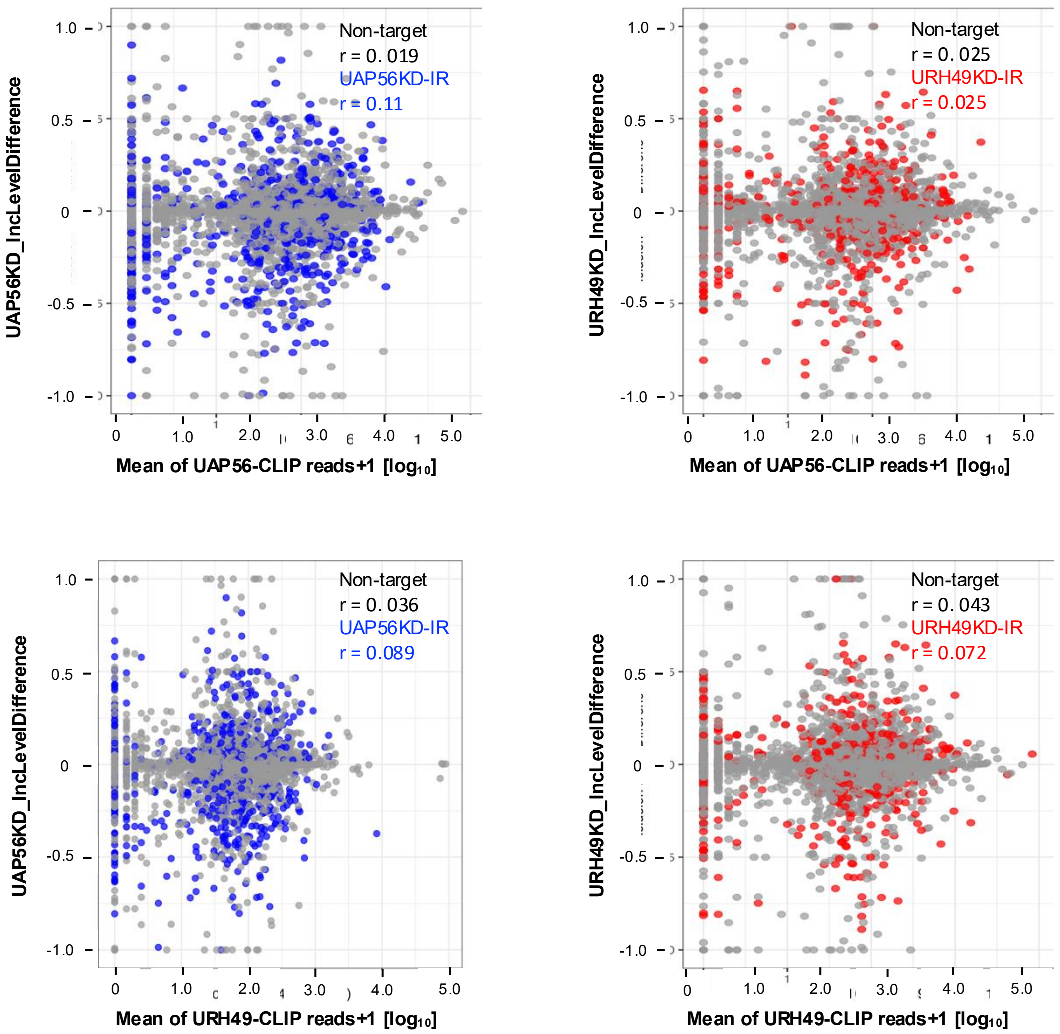
Non-target
r = 0. 019
UAP56KD-IR
r = 0.11
UAP56KD_IncLevelDifference
Mean of UAP56-CLIP reads+1 [log10]
Non-target
r = 0. 025
URH49KD-IR
r = 0.025
URH49KD_IncLevelDifference
Mean of UAP56-CLIP reads+1 [log10]
Non-target
r = 0. 036
UAP56KD-IR
r = 0.089
UAP56KD_IncLevelDifference
Mean of URH49-CLIP reads+1 [log10]
Non-target
r = 0. 043
URH49KD-IR
r = 0.072
URH49KD_IncLevelDifference
Mean of URH49-CLIP reads+1 [log10]

**Supplementary Fig.5. The non-correlation between binding and splicing regulation in the target introns and exons of UAP56 and URH49 (Related to Fig.2)**

The number of CLIP reads spanning introns and their flanking exons for UAP56 and URH49 were calculated using BEDTools IntersectBed (version 2.31.1). These values were plotted against the magnitude of the splicing changes observed at the corresponding introns upon depletion of the respective helicase. Gray dots represent introns that showed no splicing changes upon knockdown of either helicase, whereas red and blue dots indicate introns whose splicing was altered upon the depletion of UAP56 or URH49, respectively. Pearson's correlation coefficient (r) reflects the effect size.

### SFigure.6

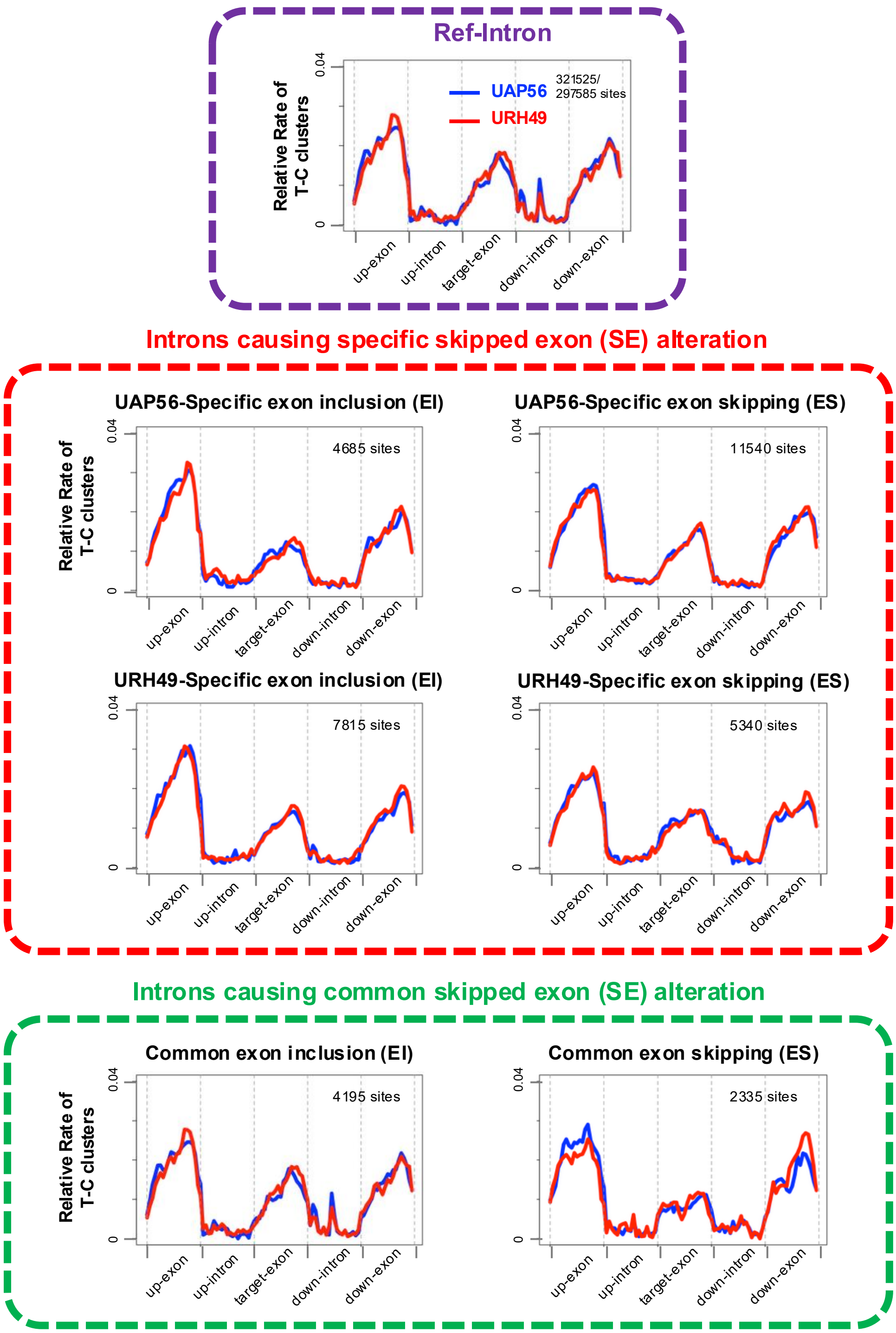

Ref-Intron
Relative Rate of T-C clusters
0.04
0
UAP56
URH49
321525/ 297585 sites
up-exon
up-intron
target-exon
down-intron
down-exon
Introns causing specific skipped exon (SE) alteration
UAP56-Specific exon inclusion (EI)
4685 sites
UAP56-Specific exon skipping (ES)
11540 sites
URH49-Specific exon inclusion (EI)
7815 sites
URH49-Specific exon skipping (ES)
5340 sites
Introns causing common skipped exon (SE) alteration
Common exon inclusion (EI)
4195 sites
Common exon skipping (ES)
2335 sites

**Supplementary Fig.6. RNA-binding patterns of UAP56 and URH49 on skipped exons (Related to Fig.2)**

Distribution of PAR-CLIP reads of UAP56 or URH49 around exon-intron boundaries. Metagene profiles show the occupancy of each binding cluster around the exon-intron boundaries. The left and right sides of each Figure represent the upstream and downstream exon-intron boundaries, respectively. The Ref-intron region covers all exons and introns.

### SFigure.7

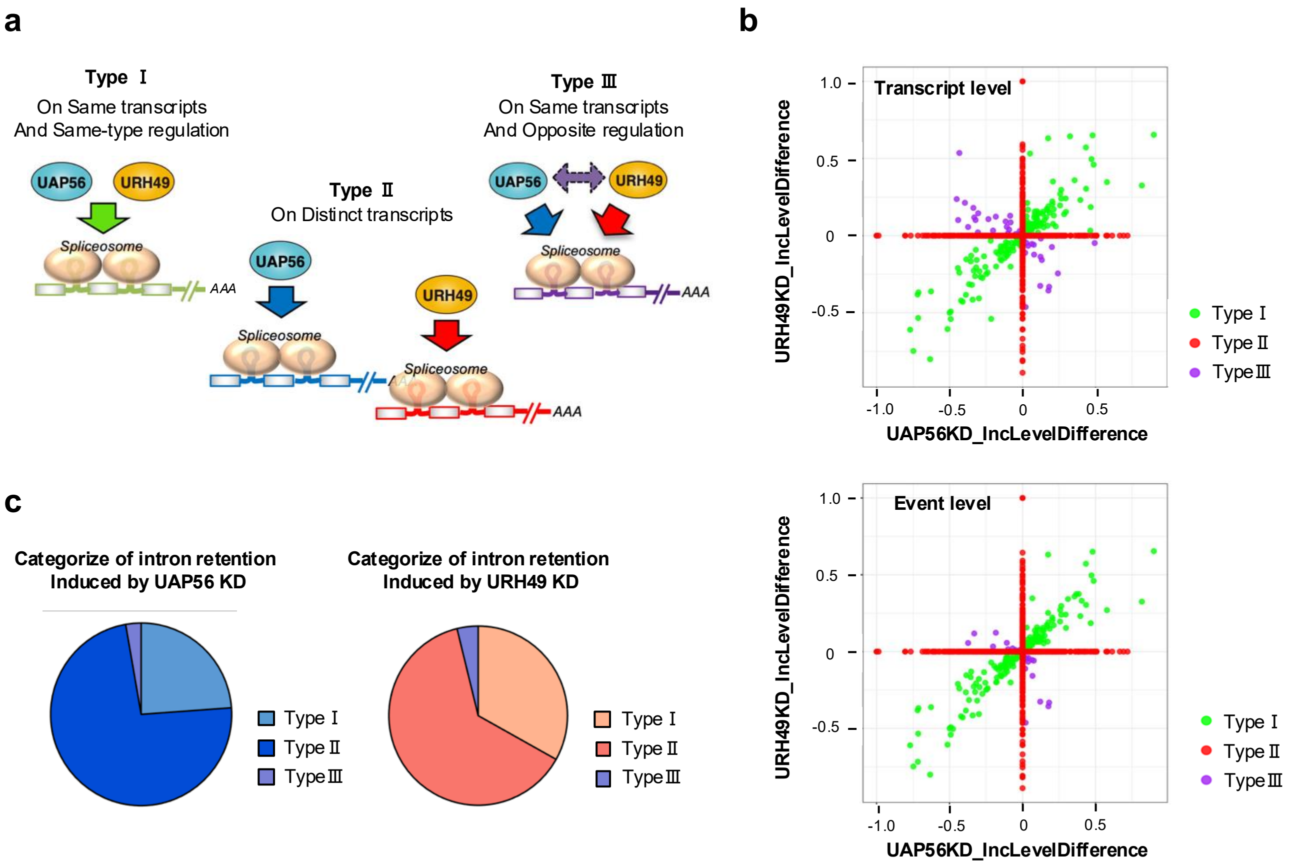

a
Type Ⅰ
On Same transcripts
And Same-type regulation
UAP56
URH49
Spliceosome
AAA
Type Ⅱ
On Distinct transcripts
UAP56
Spliceosome
URH49
Spliceosome
AAA
Type Ⅲ
On Same transcripts
And Opposite regulation
UAP56
URH49
Spliceosome
AAA
b
Transcript level
URH49KD_IncLevelDifference
UAP56KD_IncLevelDifference
1.0
0.5
0
-0.5
-1.0
-0.5
0
0.5
Type Ⅰ
Type Ⅱ
Type Ⅲ
Event level
URH49KD_IncLevelDifference
UAP56KD_IncLevelDifference
Type Ⅰ
Type Ⅱ
Type Ⅲ
c
Categorize of intron retention
Induced by UAP56 KD
Type Ⅰ
Type Ⅱ
Type Ⅲ
Categorize of intron retention
Induced by URH49 KD
Type Ⅰ
Type Ⅱ
Type Ⅲ

**Supplementary Fig.7. Three classifications of splicing regulation by UAP56 and URH49 (Related to Fig.3)**

**a** Model of the three types of splicing regulation by UAP56 and URH49. **b** Scatter plots showing transcripts from types I to III induced by the depletion of UAP56 and URH49, plotted by splicing strength (described as IncLevelDifference). The top panel shows the results of classification at the transcript level, and the bottom panel shows the results of classification at the event level. **c** Frequency of each type of splicing alteration caused by the depletion of either helicase.

### SFigure.8

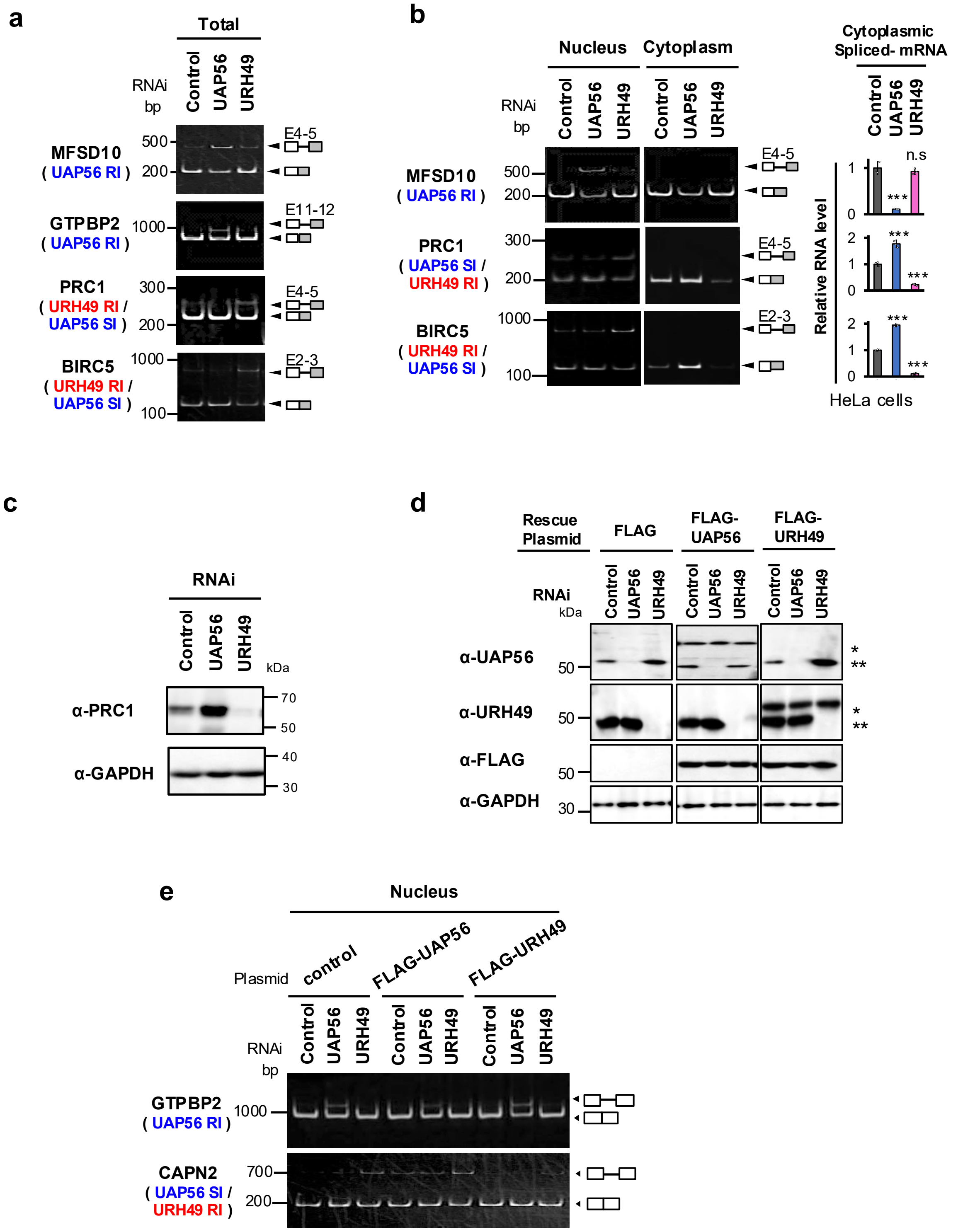
a
Total
RNAi
bp
Control
UAP56
URH49
MFSD10
( UAP56 RI )
500
200
E4-5
GTPBP2
( UAP56 RI )
1000
E11-12
PRC1
( URH49 RI /
UAP56 SI )
300
200
E4-5
BIRC5
( URH49 RI /
UAP56 SI )
1000
100
E2-3
b
Nucleus
Cytoplasm
RNAi
bp
Control
UAP56
URH49
MFSD10
( UAP56 RI )
500
200
E4-5
PRC1
( UAP56 SI /
URH49 RI )
300
200
E4-5
BIRC5
( URH49 RI /
UAP56 SI )
1000
100
E2-3
Cytoplasmic
Spliced- mRNA
Relative RNA level
n.s
***
HeLa cells
c
RNAi
Control
UAP56
URH49
kDa
α-PRC1
α-GAPDH
70
50
40
30
d
Rescue
Plasmid
FLAG
FLAG-
UAP56
FLAG-
URH49
RNAi
kDa
α-UAP56
α-URH49
α-FLAG
α-GAPDH
50
30
*
**
e
Nucleus
Plasmid
control
FLAG-UAP56
FLAG-URH49
RNAi
bp
GTPBP2
( UAP56 RI )
1000
CAPN2
( UAP56 SI /
URH49 RI )
700
200

**Supplementary Fig.8. UAP56 and URH49 contribute to mRNA export in various manners (Related to Fig.3)**

**a** UAP56 or URH49 depletion caused splicing alteration. The panels show splicing changes detected by RT-PCR using the total RNA of U2OS cells. **b** Depletion of UAP56 or URH49 caused changes in splicing and cytoplasmic mRNA expression in HeLa cells. The left panels show splicing changes detected by RT-PCR using the nuclear and cytoplasmic RNA of HeLa cells. The right panels show mature mRNA expression in the cytoplasm detected by RT-qPCR using cytoplasmic RNA. Values represent the relative expression of the indicated mRNA normalized to PGK1, as mean ±SEM of three independent experiments. Statistical analysis was performed using one-way ANOVA followed by Dunnett's test. *$p < 0.05$, **$p < 0.01$, ***$p < 0.001$. n.s.: not significant. **c** Verification of opposite changes in PRC1 expression in UAP56 or URH49 knockdown U2OS cells by immunoblotting. GAPDH was used as a loading control. **d** Confirmation of endogenous UAP56 or URH49 depletion and exogenous expression of siRNA-resistant FLAG-UAP56 or URH49 by immunoblotting. **e** Exogenous expression of FLAG-UAP56 or URH49 specifically rescued the splicing defect caused by the depletion of the corresponding factor. The panels depict splicing changes detected by RT-PCR using the nuclear RNA of U2OS cells.

### SFigure.9

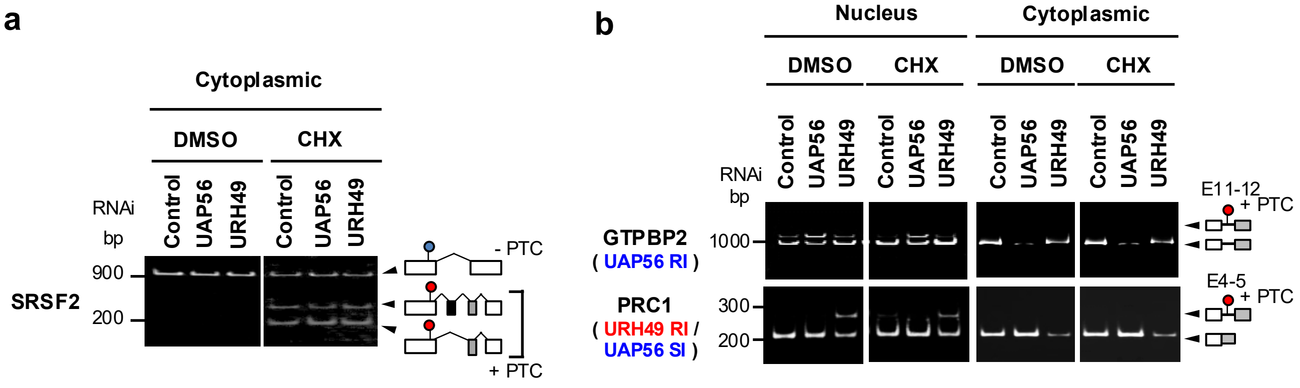
a
Cytoplasmic
DMSO
CHX
RNAi
bp
Control
UAP56
URH49
Control
UAP56
URH49
900
200
SRSF2
- PTC
+ PTC
b
Nucleus
Cytoplasmic
DMSO
CHX
DMSO
CHX
RNAi
bp
Control
UAP56
URH49
Control
UAP56
URH49
Control
UAP56
URH49
Control
UAP56
URH49
GTPBP2
( UAP56 RI )
1000
PRC1
( URH49 RI / UAP56 SI )
300
200
E11-12
+ PTC
E4-5
+ PTC

**Supplementary Fig.9. Intron-containing transcripts upon UAP56- or URH49-knockdown were retained in the nucleus (Related to Fig.3.)**

**a** The effect of nonsense-mediated decay (NMD) on mRNA with introns retained in the cytoplasmic fraction was examined. Cycloheximide (CHX) was used as an NMD inhibitor. U2OS cells were treated with CHX (50 μg/mL) for 24 h concurrently with 48 h knockdown. The RT-PCR analysis of SRSF2, a well-known target of NMD(1) was performed to verify the effectiveness of CHX-mediated NMD inhibition. Blue and red circles indicate reference stop codons and premature termination codons (PTC). + PTC indicates that the transcript variant contains a PTC. **b** RT-PCR products of the variant transcripts were detected.

1. Feng, D., Su, R.C., Zou, L., Triggs-Raine, B., Huang, S. and Xie, J. (2015) Increase of a group of PTC(+) transcripts by curcumin through inhibition of the NMD pathway. *Biochim Biophys Acta*, **1849**, 1104-1115.

### SFigure.10

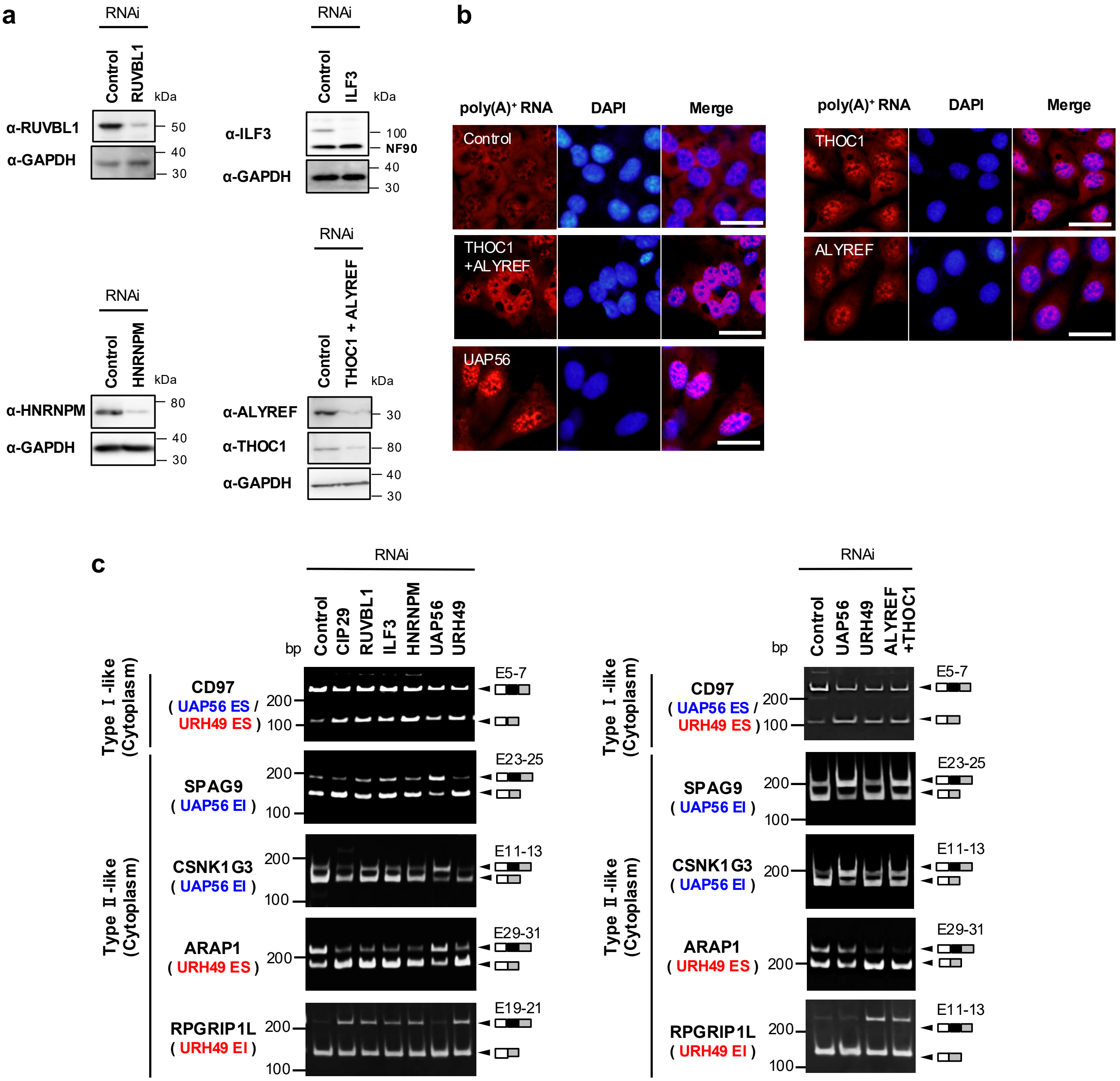

a
RNAi
Control
RUVBL1
kDa
α-RUVBL1
α-GAPDH
50
40
30
ILF3
α-ILF3
100
NF90
HNRNPM
α-HNRNPM
80
THOC1 + ALYREF
α-ALYREF
α-THOC1
b
poly(A)+ RNA
DAPI
Merge
THOC1
+ALYREF
UAP56
ALYREF
c
bp
CIP29
URH49
Type Ⅰ-like
(Cytoplasm)
Type Ⅱ-like
CD97
( UAP56 ES / URH49 ES )
SPAG9
( UAP56 EI )
CSNK1G3
ARAP1
( URH49 ES )
RPGRIP1L
( URH49 EI )
E5-7
E23-25
E11-13
E29-31
E19-21
200
100
ALYREF
+THOC1

**Supplementary Fig.10. Depletion of the apo-AREX, and the ATP-TREX complexes caused alternative splicing changes (Related to Fig.5.)**

**a** Knockdown efficiency of an individual protein in U2OS cells was validated by immunoblotting. GAPDH was used as a loading control. **b** Depletion of apo- and ATP-TREX components led to nuclear poly(A)$^{+}$ RNA accumulation in U2OS cells. DAPI was used to visualize nuclei. Scale bar, 40 µm. **c** Splicing changes were induced upon depletion of the apo-AREX or ATP-TREX complexes. The panels depict splicing changes detected by RT-PCR using the cytoplasmic RNA of U2OS cells.

### SFigure.11

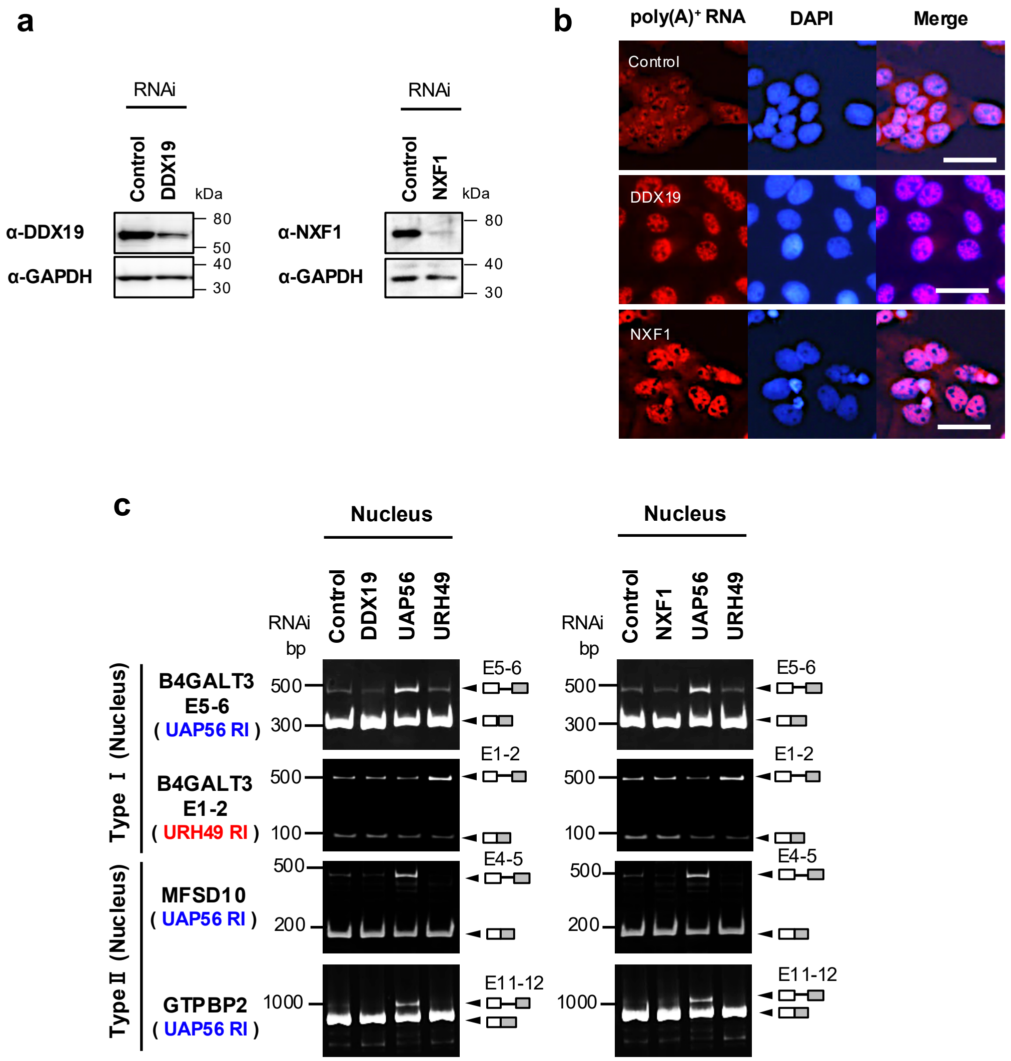

a
RNAi
Control
DDX19
kDa
α-DDX19
α-GAPDH
80
50
40
30
RNAi
Control
NXF1
kDa
α-NXF1
α-GAPDH
80
40
30
b
poly(A)+ RNA
DAPI
Merge
Control
DDX19
NXF1
c
Nucleus
RNAi
bp
Control
DDX19
UAP56
URH49
Nucleus
RNAi
bp
Control
NXF1
UAP56
URH49
Type Ⅰ (Nucleus)
B4GALT3
E5-6
( UAP56 RI )
B4GALT3
E1-2
( URH49 RI )
Type Ⅱ (Nucleus)
MFSD10
( UAP56 RI )
GTPBP2
( UAP56 RI )
500
300
500
100
500
200
1000
E5-6
E1-2
E4-5
E11-12

**Supplementary Fig.11. Depletion of the mRNA exporter does not lead to UAP56- or URH49-dependent changes in intron removal (Related to Fig.5.)**
**a** Knockdown efficiency of an individual protein in U2OS cells was validated by immunoblotting. GAPDH was used as a loading control. **b** Depletion of DDX19 and NXF1 led to nuclear poly(A)$^+$ RNA accumulation in U2OS cells. DAPI was used to visualize nuclei. Scale bar, 40 μm. **c** Depletion of the DDX19 and NXF1 did not induce splicing changes. The panels show splicing changes detected by RT-PCR using the nuclear RNA of U2OS cells.

**Supplementary Table.1 List of splicing changes detected by rMATS.**

**Supplementary Table.2 List of statistics of metagene analysis.**

**Supplementary Table.3 List of antibodies used in this study.**

**Supplementary Table.4 List of siRNAs used in this study.**

**Supplementary Table.5 List of primers used in this study.**

**Supplementary Table.6 List of RNA-seq data used in this study.**

**Supplementary Table 2**

| region | side | cor_UAP56_URH49 | p_Zdiff |
|---|---|---|---|
| Ref-Intron | ExIn | 0.9917925 | NA |
| UAP56-Specific retained intron (RI) | ExIn | 0.9657402 | 7.68E-06 |
| URH49-Specific retained intron (RI) | ExIn | 0.8840387 | 0.00E+00 |
| Common retained intron (RI) | ExIn | 0.9235385 | 2.04E-12 |
| UAP56-Specific spliced intron (SI) | ExIn | 0.9099877 | 4.02E-14 |
| URH49-Specific spliced intron (SI) | ExIn | 0.9392276 | 3.09E-10 |
| Common spliced intron (SI) | ExIn | 0.908934 | 3.02E-14 |
| Ref-Intron | InEx | 0.9923022 | NA |
| UAP56-Specific retained intron (RI) | InEx | 0.9222917 | 3.24E-13 |
| URH49-Specific retained intron (RI) | InEx | 0.8489784 | 0.00E+00 |
| Common retained intron (RI) | InEx | 0.7434267 | 0.00E+00 |
| UAP56-Specific spliced intron (SI) | InEx | 0.9294996 | 3.16E-12 |
| URH49-Specific spliced intron (SI) | InEx | 0.8660667 | 0.00E+00 |
| Common spliced intron (SI) | InEx | 0.8638791 | 0.00E+00 |
| Ref-Exons and Introns | - | 0.9963515 | NA |
| UAP56-Specific exon inclusion (EI) | - | 0.9842921 | 3.32E-07 |
| URH49-Specific exon inclusion (EI) | - | 0.9850542 | 8.23E-07 |
| Common exon inclusion (EI) | - | 0.9771103 | 1.29E-10 |
| UAP56-Specific exon skipping (ES) | - | 0.9860835 | 2.87E-06 |
| URH49-Specific exon skipping (ES) | - | 0.9832049 | 9.33E-08 |
| Common exon skipping (ES) | - | 0.9552549 | 0.00E+00 |

**Supplementary Table 3. Antibodies list in this study**

| Name | Source | Identifier |
|---|---|---|
| Goat polyclonal anti-lamin B | Santa Cruz | sc-6216 |
| Mouse polyclonal anti-GAPDH | FUJIFILM Wako | 016-25523 |
| Mouse monoclonal anti-β-tubulin | Sigma-Aldrich | T0198 |
| Mouse anti-β-actin | Sigma-Aldrich | A2066 |
| Mouse monoclonal anti-FLAG (M2) | Sigma-Aldrich | F1804 |
| Rabbit polyclonal anti-PRC1 | BioLegend | 603001 |
| Rabbit polyclonal anti-THOC1 | 10.1101/gad.1302205 | NA |
| Rabbit polyclonal anti-THOC5 | 10.1101/gad.1302205 | NA |
| Rabbit polyclonal anti-ALYREF | 10.1101/gad.1302205 | NA |
| Rabbit polyclonal anti-HNRNPM | Sigma-Aldrich | HPA024344 |
| Rabbit polyclonal anti-NXF1 | Gifted from Dr. Robin Reed | NA |
| Rabbit polyclonal anti-DDX19 | Gifted from Dr. Ichiro Taniguchi | NA |
| Rat polyclonal anti-CIP29 | 10.1091/mbc.E09-10-0913 | NA |
| Rat polyclonal anti-RUVBL1 | 10.1038/s41467-023-44217-8 | NA |
| Rat polyclonal anti-ILF3 | 10.1038/s41467-023-44217-8 | NA |
| Rat polyclonal anti-UAP56 | 10.1091/mbc.E09-10-0913 | NA |
| Rat polyclonal anti-URH49 | 10.1091/mbc.E09-10-0913 | NA |

**Supplementary Table 4. siRNA list used in this study**

| Identifier / Nucleotide sequence (d:deoxyribonucleotide ) | | | Supplier | Source |
|---|---|---|---|---|
| EGFP (Control) | sense | GGGCACAAGCUGGAGUACAACUACA | invitrogen/stealth | 10.1091/mbc.E09-10-0913 |
| UAP56 | sense | CCUGAUGAGAUAGACAUCUCCUCCU | invitrogen/stealth | 10.1091/mbc.E09-10-0913 |
| URH49 | sense | CCCUACAGCAGAUUGAGCCUGUCAA | invitrogen/stealth | 10.1091/mbc.E09-10-0913 |
| CIP29 | sense | CAAAGCCCAUUGAGCUCCCUGUCAA | invitrogen/stealth | 10.1091/mbc.E09-10-0913 |
| RUVBL1 | sense | CCACAGAGAUGGAGACCAUCUACdGdA | IDT | 10.1038/s41467-023-44217-8 |
| ILF3 | sense | GCGGAUCCGACUACAACUACGAGdAdG | IDT | 10.1038/s41467-023-44217-8 |
| HNRNPM | sense | ACAUCGAUACGAGACCUCUGAAUdTdT | IDT | 10.1038/s41467-023-44217-8 |
| THOC1 | sense | CCCUGAUAAUAUGGAAGCCUGUAAA | invitrogen/stealth | 10.1091/mbc.E09-10-0913 |
| THOC2 | sense | UGGCCAGUGGAAGAAUGAAACUUAU | invitrogen/stealth | 10.1091/mbc.E09-10-0913 |
| ALYREF | sense | GACAUGUCUCUGGACGACAUCAUUA | invitrogen/stealth | 10.1091/mbc.E09-10-0913 |
| NXF1 | sense | GAACUGGUUCAAGAUUACAAUUCCU | invitrogen/stealth | 10.1093/nar/gkac559 |
| DDX19 | sense | CAAGGUGUUUGUUCUGGAUGAGGdCdT | IDT | 10.1093/nar/gkac559 |

#### Supplementary Table 5. Primer list used in this study

| Target gene name / Primer name | | Nucleotide sequence | Aim of the primer | Source |
|---|---|---|---|---|
| UAP56 | Forward | ACTCGGTACCATGGCAGAGAACGATGTGGA | Construction of pcDNA5-FLAG-UAP56 | in this study |
| | Reverse | CCGCTCGAGCTACCGTGTCTGTTCAATGT | | |
| URH49 | Forward | CTACTCGGTACCATGGCAGAACAGGATGTG | Construction of pcDNA5-FLAG-URH49 | in this study |
| | Reverse | CCGCTCGAGTTACCGGCTCTGCTCGATGT | | |
| Halo tag | Forward | ACTCGGTACCATGGCAGAAATCGGTACTGG | Construction of pcDNA5-FLAG-Halo | in this study |
| | Reverse | TGAGGGTACCGGCGATCGCGTTATCGCTCT | | |
| P3 primer | - | CCGCTGGAAGTGACTGACAC | PAR-CLIP sample preparation | 10.1016/j.molcel.2014.05.022 |
| DSFP5 | Forward | AATGATACGGCGACCACCGACTATGGATACTTAGTCAGGGAGGACGATGCGG | | |
| DSFP3 | Reverse | CAAGCAGAAGACGGCATACGACCGCTGGAAGTGACTGACAC | | |
| SSP1 | - | CTATGGATACTTAGTCAGGGAGGACGATGCGG | | |
| tRNA | Forward | GTAGTCGTGGCCGAGTGGTTAAG | RT-PCR to confirm the fractionation | 10.1016/j.isci.2019.11.033 |
| | Reverse | GTAGTCGGCAGGATTCGAACC | | |
| U6 snRNA | Forward | GTGCTCCCTTCGGCAAGCACTTACACTAAA | | |
| | Reverse | CTCAAAAAGGAATGCTTCACAAATTTGCCT | | |
| E2F8 | Forward | CGGGGAGGAGAATAAGTACG | | |
| | Reverse | CTTGTCTTTGCGGCTGTTTA | | |
| CDK16 | Forward | GAAGCAGTACCTGGATGACTGTGG | RT-PCR to confirm alteration of intron retention events | in this study |
| | Reverse | TCTTTGTTGGGATTGACTTGGCTCG | | |
| PNKP | Forward | GCGTATGCGGAAGTCAAACC | | |
| | Reverse | GATCAAAGCCAGCCACCTTG | | |
| MFSD10 | Forward | ATTGGCTCGGCATTCTCTGTC | | |
| | Reverse | GTTCCCTTTGCTGATGCCC | | |
| GTPBP2 | Forward | TTCCATGCCACCACCTTCCGAC | | |
| | Reverse | CTGGGCCAGCACAACAAATAGCAC | | |
| DHRS1 | Forward | GAGTGGCAAATGTGTGGTGG | | |
| | Reverse | GCAAGGTCACAGGATGGCAG | | |
| CUEDC1 | Forward | CCGCGTCAACAGCCAACCTCC | | |
| | Reverse | CCTTCCTCCACCTTGGGTGCCTC | | |
| FANCI | Forward | GAGACTGCTTGATACTTGTCCTTCG | | |
| | Reverse | ACTGAGAGGATGACAGGCTGC | | |
| B4GALT3(UAP56 | Forward | GGGACTGCCTGTTCTTGCACGATG | | |

| | | | | |
|---|---|---|---|---|
| KD-RI site) | Reverse | ATTGCCCTTATCTCCTCGGTGCTTC | | |
| B4GALT3(URH49 KD-RI site) | Forward | GGCACCCAGGCACATGAAGAATAG | | |
| | Reverse | CAGAGAAAGGCTCCATCCAGCAC | | |
| PRC1 | Forward | AGTGCCTCAGTGCCCAGCTTAG | | |
| | Reverse | CATCTTCGTCTTCACACACCACATCTCTTTC | | |
| BIRC5 | Forward | CTGTAATACCAGCACTTTGGGAG | | |
| | Reverse | GAAAGGAAAGCGCAACCGGAC | | |
| CAPN2 | Forward | CGGGAGTGGCAAGCTGGG | | |
| | Reverse | TGCTTCTTCTAATGCCTTCCGC | | |
| C1ORF63 | Forward | AAGCCGTGGAATAGGTGTATC | | |
| | Reverse | ATGCTTCTTTGCTGGGTAGG | | |
| TCF7L2 | Forward | GCCAAGAGGCAAGATGGAGG | RT-PCR to confirm alteration of skipped exon events | in this study |
| | Reverse | GAGGGCTTGTCTACTCTGGAGG | | |
| SPAG9 | Forward | GGTGTCTTGTAATCGTTTGTGG | | |
| | Reverse | TGTCCCTGGAGTCACTTTATC | | |
| CSNK1G3 | Forward | AAATGCACCCATCACAGC | | |
| | Reverse | CACAAGACAGCAGATGAAC | | |
| ARAP1 | Forward | TGGCATGATGAAGTTCCGTGAG | | |
| | Reverse | GTGGGTGGCCTGAGTTTCTTC | | |
| RPGRIP1L | Forward | CCAAGACCTAAACCAAGACAAC | | |
| | Reverse | TTACCTCATCTACACTGCCTTC | | |
| METTL4 | Forward | CTCTTGGTCTGTGGAGGTAGTTGC | | |
| | Reverse | AAGTGAGCCACCACCGCTTG | | |
| CD44 | Forward | GAAAGGAGCAGCACTTCAGGAGG | | |
| | Reverse | AATGGCTTGGGTTCCACTGGGTC | | |
| CD94 | Forward | CCTGCCTGGCTTCAAGTTCATAC | | |
| | Reverse | GGTGGAGCTGTCACACTGATG | | |
| PLD1 | Forward | TCTACAAGCAGCTCCACAGG | | |
| | Reverse | GCAGTAGTCCTTTCCATGCCAG | | |
| PTS | Forward | GAAGATGAGCACGGAAGGTGG | | |
| | Reverse | ATTCATAACCATTCCCGTAGCAGG | | |
| SRSF2 | Forward | AGCGGTGTCCTCTTAAGAAAATG | RT-PCR to confirm inhibition of NMD | 10.1016/j.bbagrm.2015.04.002 |
| | Reverse | AGCATTACTCAACTGCTACACAACTG | | |
| PGK1 | Forward | GTTGCAGACAAGATCCAGCTC | qRT-PCR to confirm | 10.1091/m |

| | | | | |
|---|---|---|---|---|
| | Reverse | GAAGTGCCAATCRCCATGTTG | expression change of target gene | bc.e09-10-0913 |
| CDK16 | Forward | ATGCACAACGTGAAACTGTTCC | qRT-PCR to confirm expression change of target gene | in this study |
| | Reverse | GGTCTCGGTGTAGCACCTTC | | |
| PNKP | Forward | GCGTATGCGGAAGTCAAACC | | |
| | Reverse | GATCAAAGCCAGCCACCTTG | | |
| MFSD10 | Forward | AGTGTCCTGTTCGGAGGTCTC | | |
| | Reverse | GCTGACGTTCCCTTTGCTG | | |
| GTPBP2 | Forward | AAGGGAAAGAAGAAGAACGGAAGG | | |
| | Reverse | GTTTCCATCTTCAGCCTCGGG | | |
| DHRS1 | Forward | GACACCCTTCGCGTTGTTG | | |
| | Reverse | CAGGCTTCGCACTTCACTCTC | | |
| CUEDC1 | Forward | GTTTGTGGCGGAGTTCTGTG | | |
| | Reverse | CAGAGCAGGGGGAACCTTTG | | |
| FANCI | Forward | CCACCTTTGGTCTATCAGCTTC | | |
| | Reverse | CAACATCCAATAGCTCGTCACC | | |
| B4GALT3 | Forward | GTTGCCGTTGCTATGAACAAG | | |
| | Reverse | GGTCAGGAGTAAGTGCTGAGAC | | |
| PRC1 | Forward | GATTGAGGCAATTCGAGTGGAGCTGG | | |
| | Reverse | AGCATCGTGGAGCTGGAGCAGAC | | |
| BIRC5 | Forward | ACTGGACAGAGAAAGAGCCAAG | | |
| | Forward | AATCCATGGCAGCCAGCTG | | |
| CAPN2 | Reverse | CCCAACCTGTTCAAGATCATCC | | |
| | Forward | AGGCTTCCGTTACTTTCAACC | | |
| C1ORF63 | Reverse | CGCTACTCGCGGTCATACTC | | |
| | Forward | CCTTCCGCAGTACGACCTTC | | |
| MFSD10_Unspliced-mRNA | Forward | GTGATGCTGCTGTGCCTGGTATG | qRT-PCR to confirm alteration of intron retention events | in this study |
| | Reverse | AGGATGCGATGAGGGCCAAC | | |
| MFSD10_Spliced-mRNA | Forward | TGATGCTGCTGTGCCTGATG | | |
| | Reverse | AGAGGCTGACGTTCCCTTTG | | |
| GTPBP2_Unspliced-mRNA | Forward | CTTACTGTTCCATGCCACCACC | | |
| | Reverse | GAGAAATGGAAGGCCAGGTCCC | | |
| GTPBP2_Spliced-mRNA | Forward | CAAGGACAAACTGCGGACAGG | | |
| | Reverse | GGGCTTCTCCTGCTGTAATGG | | |
| PRC1_Unspliced- | Forward | CACTTTGGTTTTCCTCTAGGTATC | | |

| | | |
|---|---|---|
| mRNA | Reverse | GGCATCTTCGTCTTCACACAC |
| PRC1_Spliced-mRNA | Forward | AACAAAGGCTTCTAGGCGTG |
| | Reverse | TGGGGTGTGGTCTAATGCTTC |
| BIRC5_Unspliced-mRNA | Forward | GGACTGCCGCTTTAATCCCTTC |
| | Reverse | TTTGTTCTTGGCTCTTTCTCTGTC |
| BIRC5_spliced-mRNA | Forward | CCACTGAGAACGAGCCAGAC |
| | Reverse | ATGCTTTTTATGTTCCTCTATGGGG |

**Supplementary Table 6. List of RNA-seq data used in this study**

| DATA name | Source | Identifier |
|---|---|---|
| RNA-seq (Raw sequencing data) from UAP56 or URH49-depleted cells | This study | GSE280484 |
| UAP56 or URH49 PAR-CLIP-seq data | This study | PRJDB5815 |
| ALYREF and NXF1 iCLIP-seq data | 10.1016/j.molcel.2019.04.034 | GSE113953 |
